# Utilizing high resolution ribosome profiling for the global investigation of gene expression in *Chlamydomonas*

**DOI:** 10.1101/2023.02.13.528309

**Authors:** Vincent Leon Gotsmann, Michael Kien Yin Ting, Nadin Haase, Sophia Rudorf, Reimo Zoschke, Felix Willmund

## Abstract

Ribosome profiling (Ribo-seq) is a powerful method for the deep analysis of translation mechanisms and regulatory circuits during gene expression. Here, we established an optimized and high resolution Ribo-seq protocol for the unicellular model alga *Chlamydomonas reinhardtii* (Chlamydomonas). Comparing different nuclease treatments for the extraction and sequencing of ribosome-protected fragments (RPFs) and parallel RNA-seq, provided deep insight into translational dynamics and post-transcriptional control of gene expression, thoroughly covering more than 10,000 different transcripts. Our high quality Ribo-seq protocol captures the 3-nucleotide movement of elongating ribosomes along nuclear and chloroplast transcripts. Detailed analysis of the ribosomal offsets on transcripts uncovers presumable transition states during translocation of elongating ribosomes within the 5’- and 3’-sections of transcripts and features of eukaryotic translation termination. These offsets reveal drastic differences between the nature of cytosolic and chloroplast translation mechanisms. Chloroplast translation is further characterized by heterogenous RPF size distribution. We found that local accumulation of small RPFs correlates with local slowdown of *psbA* translation, possibly revealing an uncharacterized regulator step during PsbA/D1 synthesis. Further analyses of RPF distribution along specific cytosolic transcripts revealed characteristic patterns of translation elongation exemplified for the major light harvesting complex proteins, LHCs. Moreover, our Ribo-seq data can be utilized to survey coding sequence annotations and the expression preference of alternatively spliced transcripts in Chlamydomonas. We made these features easily accessible for the research community by attaching our Ribo-seq data to the most recent Chlamydomonas reference genome.

## Introduction

Translation is accomplished via ribosomes, highly conserved macromolecular ribonucleoprotein machines that decode genetic information into linear polypeptide chains resulting in an almost unlimited diversity of proteins. Protein synthesis is the origin for the quantitative and qualitative determination of a proteomés diversity, which ultimately shapes the biochemical character of a cell. It is now clear that translation output does not strictly follow the changes of transcript abundance, but instead underlies strict regulatory control (Pechmann et al., 2013; Payne, 2015; Slobodin and Dikstein, 2020). The importance of this regulation is illustrated by the fact that bacteria invest about 50% of their available energy for protein biogenesis and that more than 10% of all yeast proteins may be somehow involved in this process (Russell and Cook, 1995; Costanzo et al., 2000). Translational regulation is observed throughout an organism’s life span and includes central regulatory circuits that control the cell cycle, tissue differentiation and development, acclimation to environmental changes and the integration of external signals (reviewed in Pechmann et al., 2013; Zoschke and Bock, 2018; Teixeira and Lehmann, 2019; Wang and Amoyel, 2022). Thus, understanding the regulatory principles and mechanisms of translation is a central task of modern biology. With the tremendous advances in next generation sequencing technologies, it is now possible to systematically assay the status of cellular translation by a method called ribosome profiling (Ribo-seq) (Ingolia et al., 2009). The technology is based on the observation that ribosomes occupy precise sections of a translated mRNA (Wolin and Walter, 1988). These mRNA sections (termed ribosome footprints or Ribosome Protected Fragments, RPFs) are extracted by nucleolytic removal of un-occupied mRNA in cell lysates or isolated ribosomes. RPFs are then identified through next generation sequencing and mapping to a reference transcriptome or genome. Thus, Ribo-seq benefits from the same dynamic range as RNA-seq, and provides a rich and precise positional information of translating ribosomes from the physiological state when cells were harvested (Ingolia et al., 2009; Ingolia, 2014; Brar and Weissman, 2015). Parallel sequencing of all expressed transcripts via RNA-seq allows the comparison of RNA accumulation with translation and thereby to determine the fraction of translated transcripts, which is often referred to as “translation efficiency”. Assuming that each RPF represents a translating ribosome and thus a synthesized nascent polypeptide, mRNAs with high average RPF coverages can be considered as highly translated transcripts. In addition, local over-representation of specific RPFs within a specific protein-coding sequence (CDS) are commonly interpreted as regions of slow or even halted translation (Ingolia et al., 2009; Ingolia, 2014; Brar and Weissman, 2015). Compared to other techniques, ribosome profiling goes far beyond the targeted assays of radio-labelling of nascent polypeptides or polysome-loading for specific mRNAs, and regularly provides much deeper dataset information than proteomic analyses via mass-spectrometry (Ingolia, 2014; Brar and Weissman, 2015). Yet, ribosome profiling faces the challenges to distinguish active translation events from non-translating (paused or stalled) ribosomes. However, high-quality Ribo-seq datasets, with populations of precise read-length distribution allow the determination of active translation events if RPFs are aligned relative to the ribosomal Peptidyl-site (P-site). This reveals a 3-nucleotide periodic movement of translating ribosome along a translated CDS by over-accumulation of reads containing the first nucleotide of a decoded codon in the P-site. Thus, it is possible to distinguish ribosome pausing/stalling from translation and therefore even enables *ab initio* annotation of uncharacterized protein-coding sequences (Ingolia et al., 2009; Calviello et al., 2016; Hsu et al., 2016).

Ribo-seq has been applied to different plant species with variable gene coverage and quality, including *Chlamydomonas reinhardtii* (Chlamydomonas hereafter) (Chung et al., 2015), *Arabidopsis thaliana*, (Liu et al., 2013; Juntawong et al., 2014; Merchante et al., 2015; Hsu et al., 2016; Lukoszek et al., 2016; Chen et al., 2022), maize (Lei et al., 2015; Chotewutmontri and Barkan, 2018), and other crop plants (Wu et al., 2019; Yang et al., 2020; Yang et al., 2021; Chiu et al., 2022). Studying translational regulation in plants is of specific interest, due to the interplay of three semi-autonomous genomes (nuclear, chloroplast and mitochondrial), for which many aspects are not fully understood to date. Chloroplast gene expression seems particularly dependent on co-translational regulation, which facilitates the fast adjustment of the photosynthesis machinery to environmental changes and the stoichiometric assembly of chloroplast multi-subunit complexes which often contain nuclear and chloroplast encoded subunits (Eberhard et al., 2002; Zoschke and Bock, 2018; Fujita et al., 2019). Recently, ribosome profiling revealed the specific adjustments made to chloroplast gene expression during acclimation to different light and temperature regimes (Chotewutmontri and Barkan, 2018; Schuster et al., 2019; Chotewutmontri and Barkan, 2020; Gao et al., 2022; Trösch et al., 2022). We have previously applied a high-resolution microarray approach for the fast and cost-efficient analysis of chloroplast translation in Chlamydomonas, which allowed for the direct comparison of transcript accumulation and translation output between the alga and land plants (Trösch et al., 2018; Trösch et al., 2022). We now established Ribo-seq for the deep analyses of the three Chlamydomonas genomes. Ribo-seq had been described for Chlamydomonas before, however, with limited depth and analysis (Chung et al., 2015). However, we aimed to optimize the method to obtain a deeper coverage of genes, with super-resolution tri-nucleotide periodicity for both nucleus- and chloroplast-encoded genes and to provide a publicly accessible dataset aligned to the most recent Chlamydomonas genome (Craig et al., 2022).

## Results and Discussion

### Establishing super resolution ribosome profiling for Chlamydomonas reinhardtii

Ribosome profiling was performed from logarithmically grown Chlamydomonas cultures that were kept under mixotrophic conditions and moderate light (see Materials and Methods). Our previous experience with targeted chloroplast ribosome profiling (Trösch et al., 2018; Trösch et al., 2022) showed that addition of the elongation inhibitors chloramphenicol (CAP) and cycloheximide (CHX) (stalling 70S and 80S ribosomes, respectively) is crucial to prevent ribosome run-off during harvesting (see Materials and Methods). However, by short incubation of the inhibitors just during harvest, we reduced the exposure time to the minimum. For nucleolytic digest of RNA that is not protected by a ribosome, we applied RNase I, consistent with most Ribo-seq studies on other organisms (Ingolia et al., 2012). Our previous chloroplast ribosome profiling approaches (using MNase) revealed that the best digestion (i.e. polysome to monosome dissociation) was achieved when ribosomes were purified prior nuclease treatment (Trösch et al., 2018). We thus compared RNase I digestion on pre-purified ribosomes both at 4°C (condition i.) and 23°C (condition ii.) with samples containing whole cell lysates (condition iii.) (Figure 1A). RNase I was applied at a concentration of 1 unit (U) per micro gram (µg) RNA, which is similar to published Ribo-seq approaches performed in yeast and Arabidopsis (Chartron et al., 2016; Hsu et al., 2016; Döring et al., 2017). However, the RNase I concentration was far below the >9 U per µg RNA that was previously applied for Chlamydomonas (Chung et al., 2015). By using the lowest possible concentrations of RNase I, experimental costs can be reduced. Moreover, over digestion of ribosomes may cause an over-proportional content of contaminating rRNA fragments, which then dominate sequencing libraries. In the previous Chlamydomonas Ribo-seq study by Chung et al. (2015), contaminating rRNA fragments accounted for >90% of all sequenced reads. Our pilot runs had contaminating rRNA levels between 80 and 90%, mainly deriving from the cytosolic 5.8S rRNA and the chloroplast 23S rRNA. To further reduce the amount of contaminating rRNA species, we generated anti-sense biotinylated oligos that allow for the specific depletion of these species. For digest condition iii., we additionally tested RNase H treatment, which cleaves hybrids of the anti-sense oligos and the target rRNA contaminants. Other than the standard procedure, we performed rRNA depletion prior to dual linker ligation and were thus able to provide a higher input of RPFs in the sample, relative to contaminating rRNAs, which then allows for lowering the number of PCR amplification steps during final sequencing library generation. Importantly, all these steps were similar to a Ribo-seq protocol that we established for Arabidopsis and tobacco in parallel (Ting et al., 2023). The final PCR amplification step of the sequencing library preparation is known to be a major source of bias by over-representing certain reads (Aird et al., 2011). Read duplicates were defined as reads that share the same sequence and the same unique molecular identifier tags (UMI tags), four random bases located between a RPF and the sequencing adapters at both of its termini (Fu et al., 2018). Resulting sequencing libraries were thus corrected by deduplicating the final data sets after the genome mapping step.

**Figure 1:**
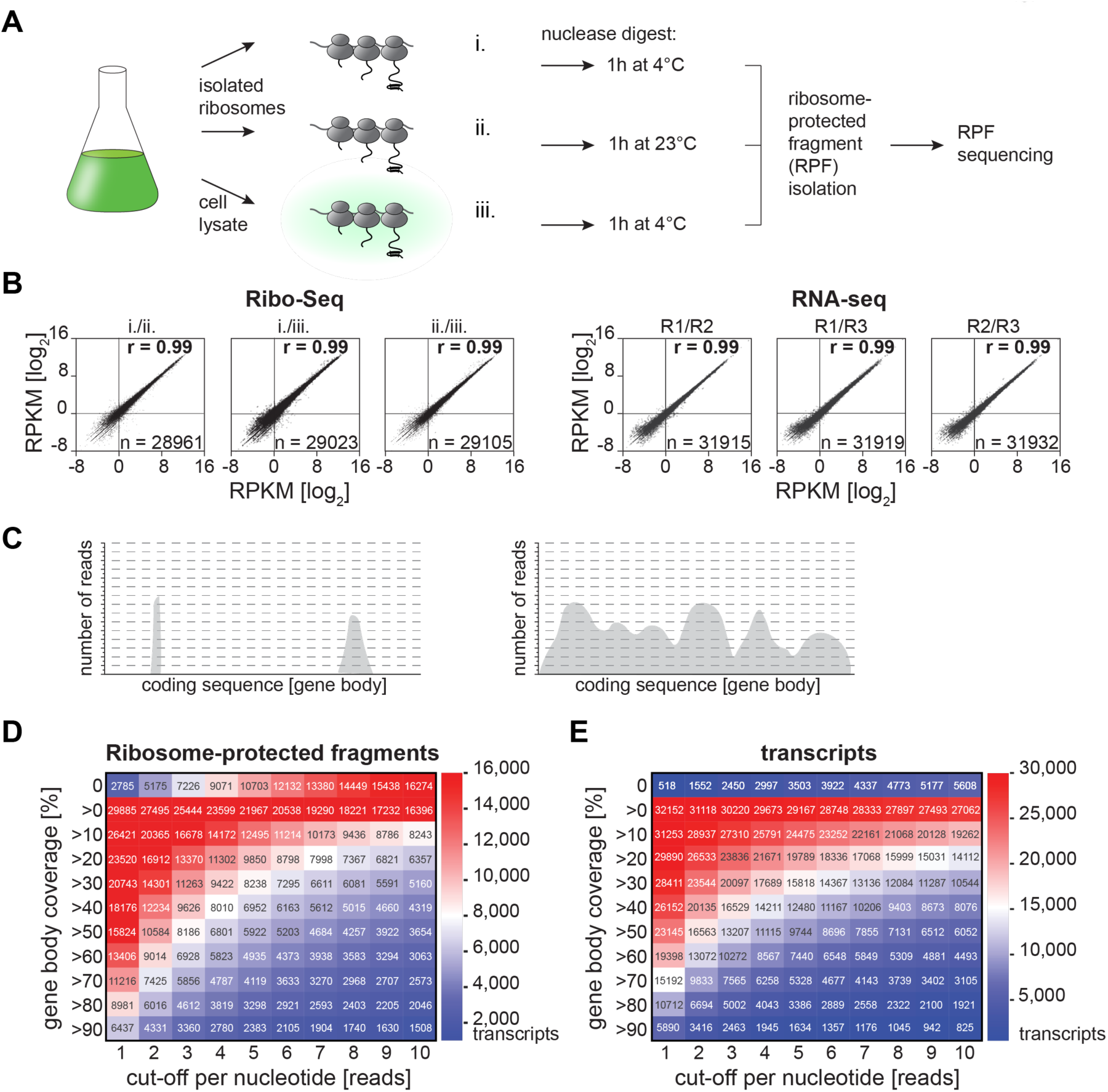
Ribo-seq provides a broad coverage of expressed genes. (A) Illustration of the experimental setup, testing nucleolytic conditions. Polysome purification and subsequent RNase I digest for 1h at 4°C (i.) or 1h at 23°C (ii.) was performed from Chlamydomonas samples, respectively. Alternatively, cell lysate was directly supplemented with RNase I (iii.). (B) Scatter plots representing the reproducibility of Ribo-seq (left panels) and RNA-seq experiments (right panels). Shown are the R-values and RPKM values. (C) Schematic representation of the concept of gene body coverage (left panel: incomplete coverage that strongly declines with an increasing read cut-off; right panel: good coverage that is robust against increasing read cut-offs). (D) and (E) Cumulative heatmaps visualizing the number of genes (given in the respective panel) covered to a certain minimal extent (rows) by a certain minimal number of reads (columns) for ribosome RPFs (C) and transcripts (D). Ribo-seq data shown are derived from the RNase I digest in lysate (condition iii.). The respective data for the other treatments are shown in Supplemental Figure 1.

Parallel to Ribo-seq, transcript accumulation was determined by sequencing whole cell RNA samples on the same platform. Ideally, RNA samples are harvested just prior to ribosome profiling experiments from the same culture. While typical transcriptomic studies enrich mRNA pools by polyadenylation-enrichment, we here sequenced fragmented small RNAs that are derived from the full set of transcripts including plastid transcripts, which frequently have no or short polyadenylation signal sequences and are thus poorly enriched by standard protocols (Komine et al., 2000; Gallaher et al., 2018).

Ribosome profiling and transcript samples were sequenced with a depth of 40 M and 20 M reads, respectively, which on average resulted in 25.7% of reads mapping to annotated CDSs of the most recent Chlamydomonas reference genome v6.1 (Craig et al., 2022). On average, 64% mapped to putative rRNA loci and 4.2% mapped to annotated untranslated regions upstream or downstream of CDSs. Despite the application of different rRNA depletion methods, the rRNA contamination was very similar in all samples. Of the sequenced reads that mapped annotated CDS a minor fraction (<1%) matched to more than 10 positions and were removed from further analyses. Of the reads mapping to CDSs, 96.24% mapped to transcripts of the nuclear genome, 3.63% mapped to chloroplast-encoded transcripts, and 0.12% to mitochondrial transcripts.

Transcript accumulation was determined by averaging reads per CDS, relative to the length of each CDS and the total read numbers of the dataset (Reads Per Kilobase per Million, RPKM), a parameter that is widely used for transcriptome analyses and accounts for a higher likelihood of measuring RNA fragments of long versus short sequences (Conesa et al., 2016). This is of particular importance for the sequencing of small and fragmented RNAs, as commonly accompanied with Ribo-seq. Averaging RPFs across the respective CDS corrects for local variations of RPF distribution and allows to get a proxy for the level of translation output per transcript (Ingolia et al., 2011; Ingolia, 2014). For better comparability and consistent with previous Ribo-seq studies, RPKM was also determined for Ribo-seq data.

Based on RPKM values, the three nuclease digestion conditions (i. to iii.) showed high reproducibility between the Ribo-seq and RNA-seq experiments, respectively (*r* = 0.99, Figure 1B). This agrees with previous observations that Ribo-seq tolerates variations in RNase digest (Gerashchenko and Gladyshev, 2017).

We next aimed to determine the depth of the Ribo-seq approach. Low coverage Ribo-seq frequently results in ribosome profiles covering only the transcripts with highest ribosome occupancy (i.e. translation output), whereas moderate or lowly translated CDS may have partial RPF coverage along the respective sequence (here termed gene body coverage, Figure 1C, left panel). For Ribo-seq with a good depth, many CDS should show decent gene body coverage (Figure 1C, right panel). The depth was determined for all Chlamydomonas transcripts of the v6 genome (32670 transcripts considering the products of gene copies). For digest condition iii., more than 10,000 transcripts had at least 50% of their CDSs sequence covered with two or more RPFs, while 6437 transcripts had over 90% of their CDS covered with at least one RPF (Figure 1D). 1508 transcripts are covered over >90% with an average of more than 10 RPFs/nt, hence representing the group of transcripts with the most complete gene body coverage combined with a high basal coverage (see also Figure 4B). Categorizing the transcripts in this way allows for the assessment of the general quality of a data set, as well as the selection of transcripts of similar quality for specific analyses without having to rely on expression values alone. The gene body coverage was highest for digest condition iii. (Figure 1D and Supplemental Figure S1), and comparably good for the coverage of RNA fragments for the RNA-seq experiment (Figure 1E).

The read length distribution of the RPFs mapping to annotated CDSs of the nuclear genome displayed high comparability to published Ribo-seq approaches with other organisms. RNase I treatment typically results in RPFs with a length between 28 and 30 nt. Here, digest of purified ribosomes at 4°C resulted in RPFs with a predominant length of 31 nt. Digest condition ii. caused a broader distribution of RPFs between 28 and 30 nt, while digest in cell lysate resulted in a sharp peak of RPFs with a length of 30 nt (Figure 2A). Such a sharp peak at 30 nt is typically found in Ribo-seq from mammalian cells when harvested in the presence of CHX (Wu et al., 2019; Sharma et al., 2021). The sharp peak is even more prominent if only those RPFs are considered that map to the CDS (Supplemental Figure S2A). This is most evident for the size distribution mapping to the CDS of the 8 mitochondrial transcripts. Here, RPF accumulations peaks at a size of 33 nt (Supplemental Figure S2A). The larger size of mitochondrial RPFs was described before and could stem from the bulkiness of the membrane associated ribosomes (Rooijers et al., 2013). Whereas, cytosolic and mitochondrial ribosomes produced sharp RPF peaks, the RPF size distribution of chloroplast transcripts is less defined and may be the result of diverse translational states, as previously reported for maize and Arabidopsis (Chotewutmontri and Barkan, 2016; Gawronski et al., 2018; Fujita et al., 2019) (see below). The sharpest peak for chloroplast RPFs is seen under condition ii., albeit the predominant RPFs of 28 nt is found under all treatment conditions (Figure 2A). While consistent with the observations in other organisms, our RPF size distribution varies from the previous Ribo-seq data with Chlamydomonas, which reported a general shift towards smaller RPF sizes, maybe as consequence from their harsh RNase I treatment (Chung et al., 2015).

**Figure 2:**
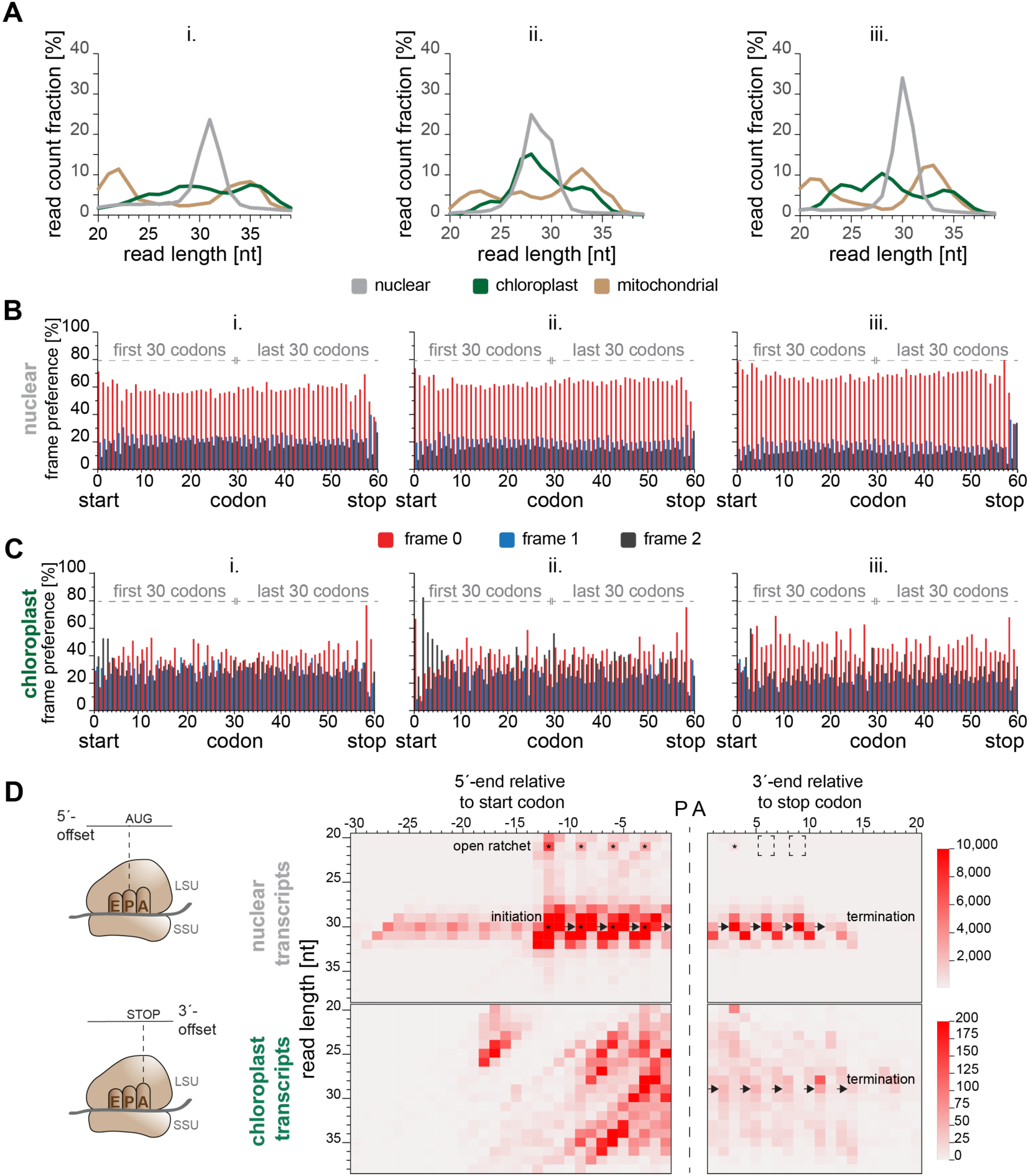
Tri-nucleotide periodicity of cellular translation. (A) Length distribution of RPFs for each organelle from the three different nuclease treatments tested (i. and ii.: digest of purified ribosomes at 4°C and 23 °C, respectively, iii.: digest in lysate at 4°C). (B) and (C) metagene relative frame preference of nuclear and chloroplast-encoded transcripts per codon for all three nuclease treatments. Codons 1-30 represent the first 30 codons of each transcript while codons 31-60 represent the last 30 codons of each transcript. Frame 0 corresponds to AUG in the ribosomal P-site. (D) Left side: cartoon of the ribosome with small (SSU) and large (LSU) subunits. Heatmaps representing normalized 5’-P-site offset and 3’-A-site offset counts for nucleus- and chloroplast-mapped RPFs that fully enclose a start or stop codon. Columns represent 5’/3’-RPF end positions relative to the start or stop codon while rows represent the RPF lengths. The shades of red are proportional to the number of RPF ends mapping onto a tile. Arrows indicate the triplet-wise movement of ribosomes in early or late elongation state. Asterisks indicate the tiles harboring the most 5‘- or 3‘-ends of the 21 and 30 nt RPF-species. Dashed boxes indicate the sudden absence of 21 nt RPFs for the last two codons before translation terminates.

A hallmark of high-resolution Ribo-seq data is the triplet-wise 3-nt periodicity of RPFs that represents slowdown of ribosome movement as ribosomes decode a respective codon (Ingolia, 2016). To calculate periodicity, it is necessary to determine the 5’-offset for every RPF length species, which is the number of bases located 5’ to the P-site (the translational state of polypeptide bond formation) of a respective RPF. This is usually done by considering all RPFs that fully enclose an annotated start codon and determining the frequency of all occurring 5’-offsets for every RPF length separately. The 5’-offset, most frequently occurring for a specific RPF length, is then declared to be the true 5’-offset of this species. This strategy is based on the fact that RPF occupancy of transcripts usually inclines dramatically at the borders between 5’-untranslated regions and CDS, with the start codon often having an exceptionally high coverage, which is due to the slow process of translation initiation (Supplemental Figure S2B). The 5’-offsets determined in this way are used to calculate the exact P-site position and reading frame of every RPF. By this, 3-nt periodicity can be nicely visualized in meta-plots combining all genes while 3-nt plots of individual transcripts can be noisy, depending on translation levels. The exact 5’-offset varies in an organism- and protocol-specific manner and is usually 12 to 13 nt. Alternatively, the P-site position has also been determined by calculating the 3’-offset of the ribosomal Aminoacyl-site (A-site), applying the same strategy to RPFs mapping to annotated stop codons.

With the majority of cytosolic RPFs with one read length (Figure 2A), all three RNase I digest conditions yielded a clear frame preference of “frame 0” for nucleus-encoded transcripts, meaning that the base triplet found at the P-site of a RPF is for the for the majority of RPFs in-frame with the annotated CDS. Again, lysate digest (iii.) resulted in the best frame preference values (>68% of RPFs in frame 0, 19.4% in frame 1 and 12.5% in frame 2), suggesting that this condition yields Ribo-seq data of the best quality (Figure 2B and Supplemental Figure S2B). Due to the differences between 80S and 70S ribosomes, 3-nt periodicity needs to be determined separately for chloroplast translation. For chloroplast transcripts, the frame preference was less obvious, albeit it was clearest for digest condition iii. (Figure 2C and Supplemental Figure S2B). Considering the overall quality of sample iii. (the similar degree of rRNA contamination between all three samples and the high correlation with the other samples on RPKM level), it can be concluded that RNase H mediated rRNA depletion has no adverse effects on the Ribo-seq sample quality and seems to be similarly effective.

To get a better insight into the RPF offsets and periodicity, different lengths of the RPFs were plotted against their offset frequencies upon P-site and A-site alignment for the dataset of condition iii. (Figure 2D). The triplet-wise movements of the 5’ends with an offset of 12 nt during initiation is clearly visible for cytosolic RPFs and most prominent for the 30 nt RPFs of digestion conditions iii., consistent with the observation in other eukaryotic organisms (Lauria et al., 2018). Likewise, triplet-wise movements of the 3’-ends are also detectable and are in frame with the stop codon. Here, a 9 nt long 3’-offset seems the predominant form prior to termination with the stop codon in the A-site of the ribosome. However, due to the 12 nt 5’-offset and 3 nt occupied by the P- and A-sites respectively, a 3’-offset of 12 nt was expected. This discrepancy could be a result of CHX treatment, which locks ribosomes in a pre-translocation conformation after peptide bond formation (Wu et al., 2019) so that RPFs on non-stop codons may exceed the natural termination-caused accumulation on stop-codons. Moreover, a slow termination process might arrest ribosomes on the codon just upstream of the stop, prolonging ribosome dwell time at this site. At the same time, ribosomes that already entered termination during sampling are likely to complete the process and dissociate from the transcript. CHX does not inhibit termination, which in turn would dramatically lower the number of FPs mapped to stop codons with their A-site. Interestingly, RPFs with a length of 21 nt are detectable upon initiation (asterisks, Figure 2D). These fragments have been previously characterized as RPFs of ribosomes in an open ratchet conformation during translocation, even accumulating as abundant species without CHX treatment in yeast (Lareau et al., 2014). During elongation, ribosomes undergo massive conformational changes by rotating the large subunit relative to the small subunit (Frank and Agrawal, 2000; Zhang et al., 2009). The fact that the RPFs produced from the open-ratchet conformation have the same 5’-offsets as the main RPF species despite being considerably shorter, suggests that the RNase exclusively removes nucleotides from the 3’-end of RPFs in this conformation (and could indicate that the center of the rotation is located towards the 5’-direction, viewed relative to the RPF). Cycloheximide was shown to prevent binding of the next aminoacylated tRNA in the A-site during peptide bond formation (Schneider-Poetsch et al., 2010) and thus stabilizes ribosomes in a specific stage during the elongation cycle, in which RPFs of 28-30 nt are the prevalent form. Interestingly, human 80S ribosomes are not exclusively arrested in one conformation upon CHX treatment (Sharma et al., 2021), comparable to the situation seen here for Chlamydomonas 80S ribosomes (Figure 2D, upper panel). These 20-22 nt RPFs might derive from a post-translocation ribosome species (Lareau et al., 2014). The presence of both species demonstrates active movement of ribosomes along the CDS. The absence of the “open-ratchet” 21 nt RPFs starting from two peptide-bond formations prior to termination, may support the idea from above that ribosomes near stop codons are paused, which can prevent the collision between ribosomes (Figure 2D upper panel, dashed box). Interestingly, the frame preference for frame 0 of the stop-minus-2 codon is the highest in the whole metagene periodicity plot (79.6% Figure 2B), which could denote a short stalling at this codon before translocation. However, for the next codons the frame 0 preference drops considerably to 55.8%, especially in favor of frame 1 (36.2%) and then to 33.1% at the stop codon, clearly indicating that translation dynamics change towards the termination step. Moreover, these characteristics are accompanied by a continuous decline of absolute numbers of reads mapping to these codons on the meta-gene level (for digest iii: stop-2: 19890, stop-1: 6786, stop: 522).

The prokaryotic-type chloroplast RPFs 5’-offsets seem to be much more diverse than their cytosolic counterparts, apparent by their multiple clusters on the heat map (Figure 2D, bottom panel). These might resemble populations of initiation complexes in different states or in complex with transcript specific RNA processing factors and *trans*-acting factors that interact with ribosomes (Westrich et al., 2021). Without clear criteria how to distinguish and interpret these clusters, 5’-offset definition is problematic, since multiple 5’-offset frequencies of comparable amplitude exist for many RPF lengths. However, the chloroplast 3’-offsets seem less scattered and reveals a main RPF species of 28 nt length with 11 nt 3’-offset. This is in accordance with the RPF length distribution of all chloroplast RPFs, where the 28 nt species also represents the main species, although the distribution is much broader than for the cytosolic RPFs. Similar to the cytosolic offset heat map, a triplet-wise movement of ribosomes can also be observed for the terminating chloroplast ribosomes in the 3’-offset heatmap for 29 nt RPFs, albeit with much lower clarity. Altogether, the observations made for chloroplast offsets and RPF length distribution might indicate a certain heterogeneity of the chloroplast ribosome pool, which is an interpretation that would be in line with the large number of proteins that interact with chloroplast ribosome found in our previous study (Westrich et al., 2021) and the large number of RNA-binding proteins known or suspected to regulate the translation of specific chloroplasts transcripts (Nickelsen et al., 2014). This is remarkable considering the very small number of transcripts encoded on the chloroplast genome (72). Together, this might suggest that translation in the chloroplast of Chlamydomonas is heavily influenced by transcript-specific features rather than following a universal scheme (Zoschke and Bock, 2018).

### Differences in translation output between cytosolic and organellar transcripts

We next compared the translation levels of the organellar versus nuclear transcripts. Chlamydomonas cells contain on average 83 plastid genomes within the large cup-shaped chloroplast occupying about 40% of the cellular volume. The mitochondrial genomes exist in 130 copies, also greatly outnumbering the one haploid nuclear genome (Gallaher et al., 2018). Despite the rather low contribution of organellar RPFs relative to the total RPF pool, both chloroplast and mitochondrial transcripts had higher RPKM values when compared to the average RPKM values of nuclear genes (Figure 3A). A similar trend is seen for transcript levels (Figure 3B), suggesting that organellar genes are expressed at considerably higher levels compared to most nuclear genes (Forsythe et al., 2022).

**Figure 3:**
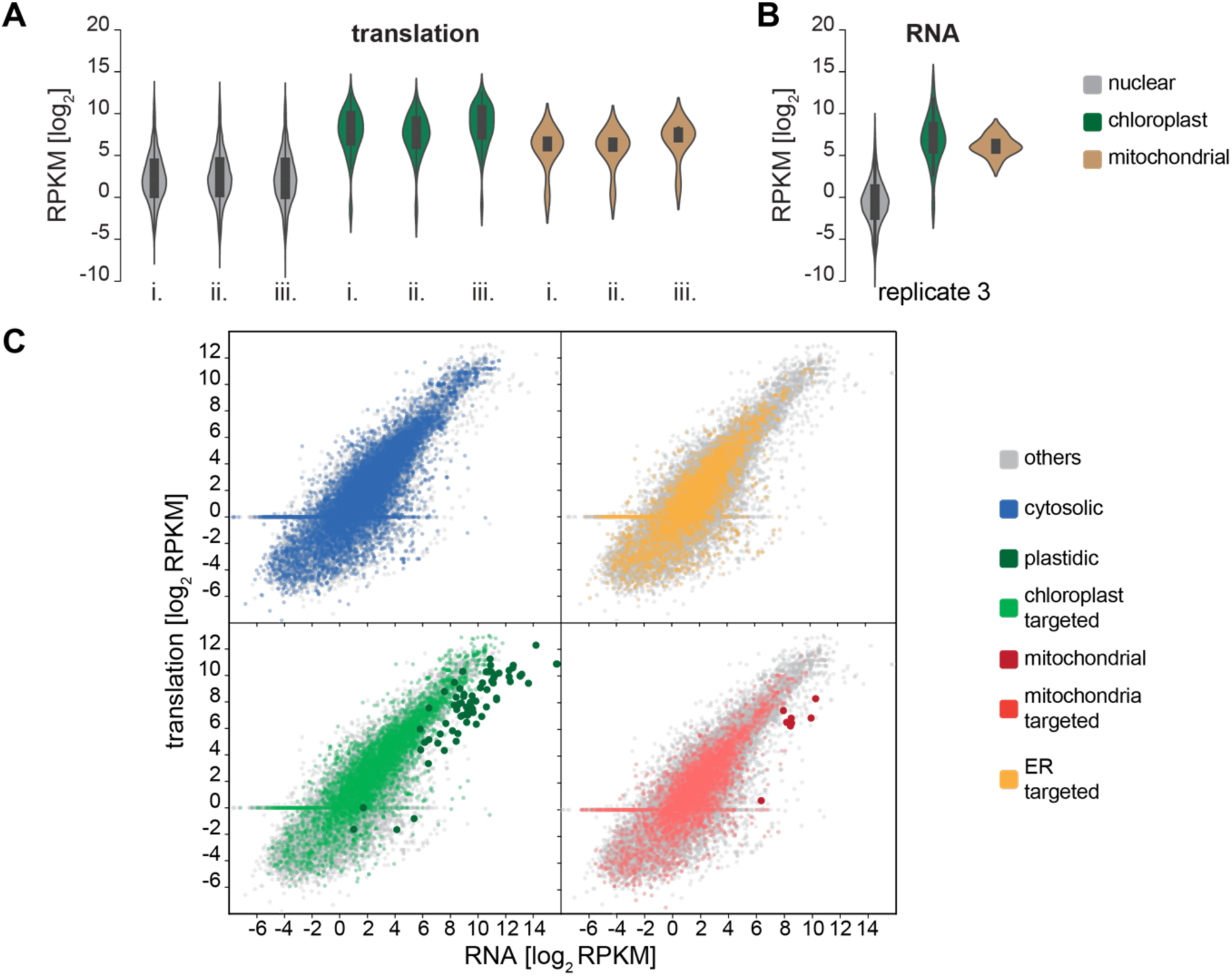
Chloroplast-encoded proteins are highly expressed. (A) Violin plots of the RPF data, displaying the RPKM (count per million reads) distribution for genes of each plant cell genome, following the three different RNase I treatments (i. and ii.: digest of purified ribosomes at 4°C and 23°C, respectively, iii.: digest in lysate at 4°C). Values with a RPKM cut-off <2 (log2) are excluded. (B) Violin plots, displaying distribution of RPKM (reads per kilobase of exon per million reads mapped) for RNA values with a cut-off >2 (log2) for each plant cell genome in replicate 3 exemplary. (C) Scatterplots of the averaged Ribo-seq and RNA-seq (RPKM) data for different subcellular-localized proteins. Coloring indicates subcellular localization and origin of the encoded proteins.

With the high reproducibility of Ribo-seq and RNA-seq data, we chose to average the datasets for estimating the levels of transcript accumulation and translation output. A direct comparison between translation output and transcript accumulation is frequently referred to as translation efficiency (TE) and normalizes RPF values over transcript accumulation. Contrasting translation over transcript data of the Chlamydomonas genes in a scatter plot and highlighting their subcellular localization showed that nuclear transcripts encoding chloroplast-localized proteins have higher expression values when compared to proteins of cytosolic or Endoplasmic Reticulum localization (Figure 3C), again supporting the notion that many chloroplast proteins are expressed at high levels. Interestingly, most nucleus-encoded transcripts showed similar levels between transcript accumulation and translation (distributed along the diagonal, Figure 3C). In contrast, chloroplast- and mitochondria-encoded transcripts showed higher transcript accumulation levels, when compared to translation, which agrees with the published data of profound potential translational regulation within the chloroplast (Sun and Zerges, 2015; Trösch et al., 2018; Zoschke and Bock, 2018). Remarkably, translational regulation seems also predominant for the few mitochondrial genes.

Column plots of the individual chloroplast transcript and translation values show that even the weakest expressed known chloroplast transcripts have RPKM values ∼10, which is within the median range of nucleus-encoded transcripts (Figure 4A versus 3A). Not surprisingly, transcripts that encode the core photosystem I subunits PsaA and PsaB, the photosystem II core subunits PsbA/D1,PsbB/CP47, PsbC/CP43, PsbD/D2, the Rubisco large subunit, RbcL, the CF_0_ ATPase subunit AtpH, and the translation elongation factor TufA show the highest translation levels, consistent with our previous chloroplast ribosome profiling experiments and the examination of protein accumulation levels (Schroda et al., 2015; Trösch et al., 2018). We further detected transcripts of the uncharacterized chloroplast Open Reading Frames (ORF) *orf202* (positioned within the transposon Wendy I), *orf854* (positioned within the transposon Wendy II), *orf528*, and *i-crel* (Gallaher et al., 2018), however, translation output was low or even undetectable (for *orf2020* and *i-crel*), suggesting that these transcripts are barely translated in Chlamydomonas cultures, at least under standard conditions (Figure 4A). The direct TE ratios of the chloroplast transcripts again show the imbalance between chloroplast transcript accumulation and translation output, suggesting that the long-living chloroplast transcripts present a buffering capacity for the fast and efficient adjustment of translation levels during environmental changes, as for example seen during light and temperature acclimation and most prominent for PsbA/D1 synthesis (Chotewutmontri and Barkan, 2018; Schuster et al., 2019; Trösch et al., 2022).

**Figure 4:**
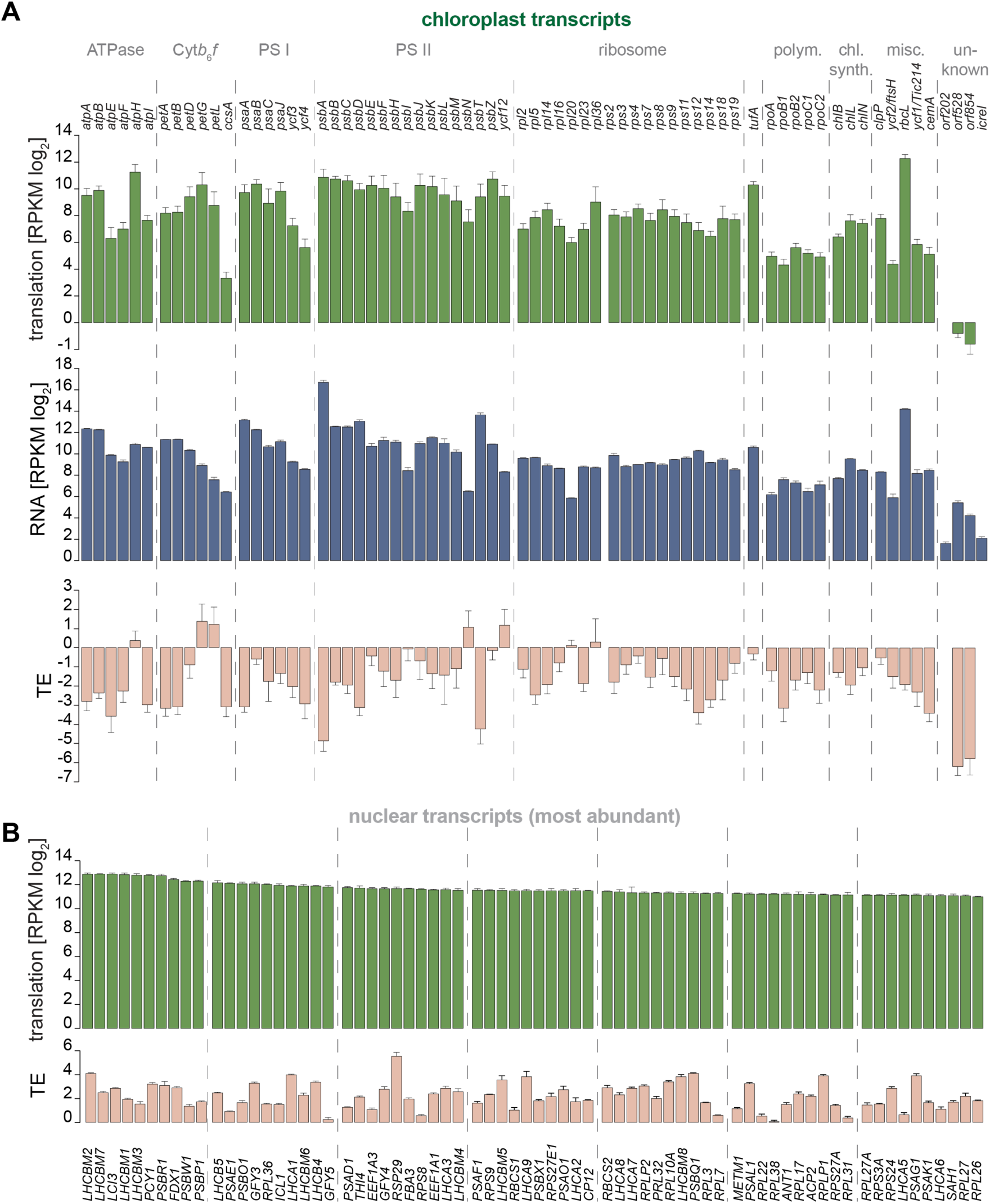
Translation output of chloroplast- and nucleus-encoded transcripts. (A) Mean read intensity/TE ratio (per respective coding sequence) of chloroplast-encoded transcripts on the level of RPF, RNA and translation efficiency (TE). The transcripts are grouped by function of the encoded proteins (Cyt*b*6*f*= cytochrome *b*6*f* complex, PS I = photosystem I, PS II = photosystem II, polym. = RNA polymerase, chl. synth. = chlorophyll synthesis, misc. = miscellaneous). (B) Values of the 70 most highly expressed nucleus-encoded genes on the translation level. Only the longest splice variants were considered. Respective translation efficiency for these transcripts is given below. For comparison, transcripts with the top 70 highest TE values are given in Supplementary Figure S3C. Error bars indicate standard deviation of the three experiments (treatment i. to iii. for Ribo-seq, and biological replicate 1-3 for RNA-seq), and are shown unidirectional to improve readability.

We next determined translation output levels of nucleus-encoded transcripts, only considering the longest transcript variant if multiple splice forms existed. Of the transcripts with the highest translation output, 51 protein products have a known or predicted chloroplast localization and 47 are present in the nucleus or cytosol, again showing the high accumulation levels of chloroplast proteins (Supplemental Figure S3A). Amongst those, transcripts of the light harvesting complex proteins (LHCs) have the highest translation output, consistent with their high protein accumulation levels in plant cells (Dall’Osto et al., 2015). Generally, *LHC* translation showed comparable RPKM translation output values, whereas transcript accumulation was rather variable. For the transcripts of the high-light induced photoprotective LHCSR3.1 and LHCSR3.3 proteins, we could not determine translation profiles (Supplemental Figure S3B), suggesting that these proteins are not synthesized under the investigated growth conditions (Bonente et al., 2012). Besides photosynthesis related transcripts, the list with top-most translation contained transcripts encoding numerous ribosomal proteins of the 80S complex and elongation factors (Figure 4B). Other abundantly translated proteins are the putative acetate transporter GYF3 (RPKM 4285, rank 14), the Isocitrate lyase (ICL1, RPKM 3918, rank 16), a key enzyme of the glyoxylate cycle (Plancke et al., 2014), the thiazole biosynthesis enzyme (THI4, RPKM 3402, rank 22) (Moulin et al., 2013) or the Methionine Adenosyl-transferase (METM1, RPKM 2438, rank 51), confirming their central role in Chlamydomonas cells. Plotting of transcripts with the highest TE values showed an over representation of histone transcripts, suggesting their robust translation in Chlamydomonas cells under logarithmic growth (Supplemental Figure S3C).

### Exploiting individual ribosome profiles to mine for putative regulatory principles

Local elongation rates are known to vary considerably during translation along a transcript, depending on tRNA availability, mRNA secondary structure, the amino acid sequence of the resulting polypeptide and co-translational folding (Gloge et al., 2014; Kramer et al., 2019). With the robust gene body coverage over Chlamydomonas transcripts (Figure 1D), we aimed to learn more from the dynamics of translation elongation, derived from the RPF distribution over individual coding sequences (ribosome profiles). Ribosome profiles were highly comparable between the three RNase I treatment conditions i.-iii., as indicated by median *Pearson* correlations between data sets ii. and iii. of ∼0.62 for nucleus-encoded transcripts and >0.85 for chloroplast-encoded transcripts. Although RNA-seq read distributions over individual CDS were even more comparable (median of *r*-value 0.87 for nucleus-encoded and 0.99 for chloroplast-encoded transcripts) among the RNA-seq data sets, they showed no correlation (median *r*-value close to zero) to the respective ribosome profiles (Figure 5A and Supplemental Figure S4). Certainly, methodological-driven differences between Ribo-seq and RNA-seq cannot be fully excluded, but it can be assumed that the ribosome profiles reflect transcript-specific translation dynamics and are likely not the result of sequencing biases.

**Figure 5:**
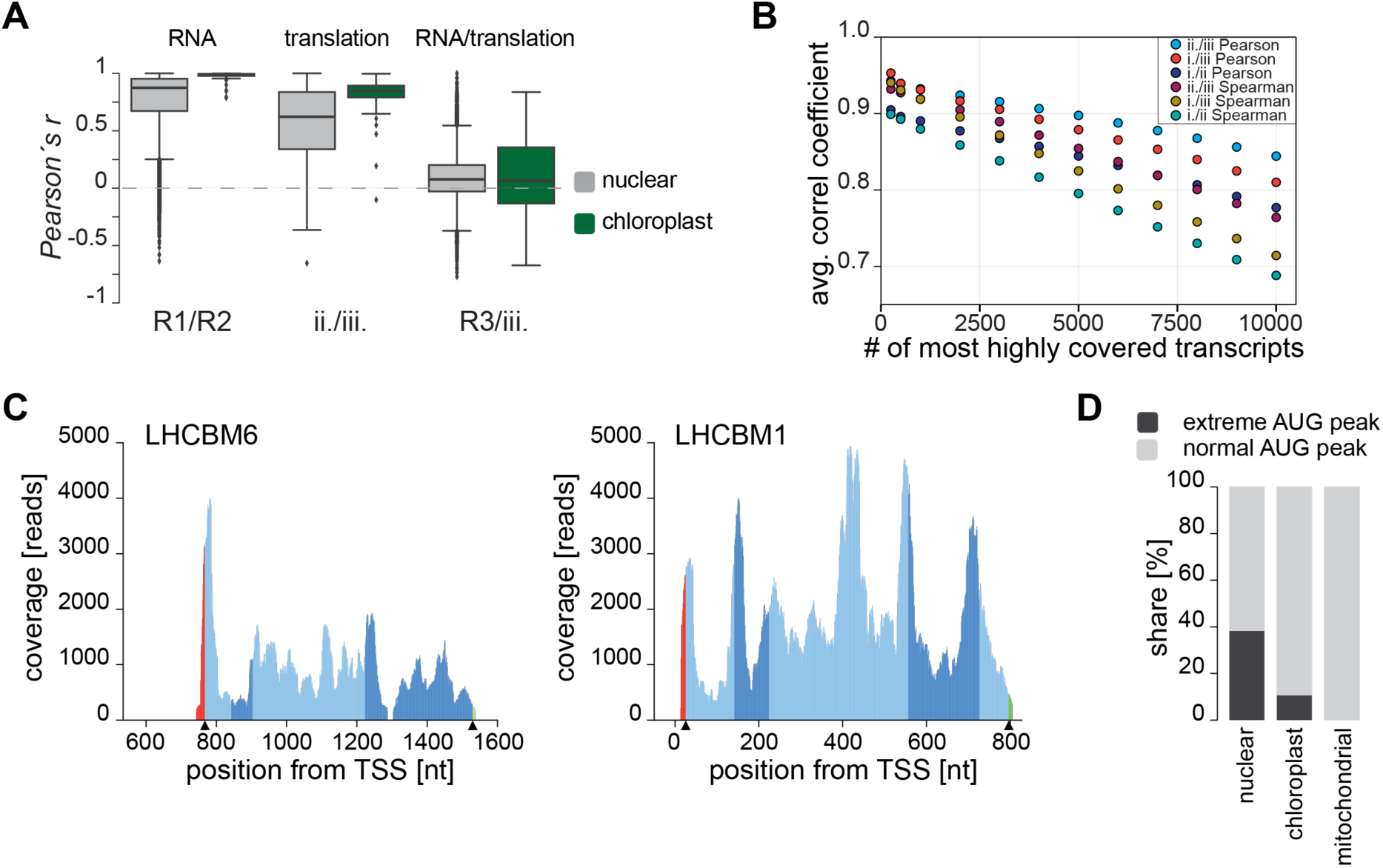
Putative translational regulation of individual transcripts. (A) Boxplots representing the distribution of Pearson’s *r* correlation coefficients comparing the RNA gene body coverage and ribosome profiles of all chloroplast- and nucleus-encoded transcripts between replicates (R1 and R2 for RNA) and between two types of RNase I treatments (ii. and iii. for translation). (B) Average *Pearson* and *Spearman* correlation coefficients depending on the number n of considered transcripts. Transcripts were ordered by descending RPF count per transcript, n most highly covered transcripts were selected, correlation coefficients were computed for each transcript in this subset, and the average was plotted. For correlation coefficients of individual transcripts see Supplemental Figure S5A. (C) Ribosome profiles of LHCBM1 exemplary for a profile with even RPF distribution and LHCBM6 as an example for a profile with an extreme initiation peak. Red parts of the profile represent coverage within the 5’-UTRs, green parts represent 3’-UTRs and alternating blue shades represent exons within the CDS. Black arrows on the x-axis indicate start and stop codons. (D) Stacked column chart representing the fraction of ribosome profiles containing extreme initiation peaks (defined as transcripts with an average coverage in first 7 codons exceeding by at least 4 times the average coverage of the whole CDS). Nucleus-, chloroplast- and mitochondria-encoded transcripts are plotted separately containing only transcripts with a gene body coverage of >70% in the Ribo-seq (transcripts with extreme initiation peak are 3346 out of 8757 for nuclear, 7 out of 66 for chloroplast and 0 out of 7 for mitochondria). Further information on occurrence of abnormal initiation peaks is provided in Supplemental Figure S6.

The correlation coefficients for specific transcripts comparing two experiments (both *Pearson* and *Spearman*) stably range between 0.7 and 1 for the majority of ribosome profiles that have an average coverage >100 reads per nucleotide, suggesting that low correlation coefficients are the consequence of sequencing depth limitations and thus limited observability of lowly translated transcripts rather than the true absence of a characteristic translation dynamic of these transcripts (Supplemental Figure S5). Importantly, more than 10,000 transcripts have *r*-values >0.7 (Figure 5B), again showing the high reproducibility between the RNase I digest conditions for transcripts with moderate and high RPF coverage (Supplemental Figure S5). In fact, the *r*-value distribution shows that these direct comparisons of read coverage per transcript might be well suited for selecting good cut-off ranges for a deeper analyses of ribosome profiles. In many previous studies, simply minimum RPKM values (e.g. 1) were taken as cut-off.

Ribosome profiles of the highly translated *LHC* transcripts showed some remarkable differences between the individual *LHC* species. Efficiently controlling the expression of *LHC* genes is essential for regulating the antenna size of the photosynthesis machinery in order to capture photons under low irradiance conditions (increased expression) or avoid over-excitation and thus photodamage under high irradiance conditions (reduced expression) (Dall’Osto et al., 2015). Of the transcripts encoding the 21 different LHC members serving Photosystem I or II (LHCA and LHCB, respectively), 20 were found amongst the top 100 highest translated transcripts (not considering splice variants). Only the transcript encoding LHCB7, a rather newly discovered protein containing the unusual number of 4 transmembrane domains (Klimmek et al., 2006) had low RPKM values of ∼15 (Supplemental Figure S3B). The variations in *LHC* transcript levels result in considerably different TE values (Supplemental Figure S3B), suggesting that translational regulation might be achieved by different strategies for expression of the *LHC* species. Indeed, the ribosome profiles are considerably different between transcripts encoding the different LHCs. For example, *LHCBM6* showed an exceptionally strong peak of RPFs covering the first 7 codons, which exceeds the average RPF density over the remaining CDS by more than 4-fold (Figure 5C). While “initiation peaks” are commonly accumulating over the first few codons of a CDS, their amplitudes are usually within the dynamic range of the remaining CDS. Such high peaks, in contrast, could point to a regulative mechanism stalling ribosomes during or shortly after initiation. Indeed, *LHCBM6* was shown to be translationally repressed under moderate light conditions by a co-translational regulator termed NAB1 (Nucleic Acid Binding protein). NAB1 itself is tuned by a regulatory circuit sensing carbon dioxide supply levels in Chlamydomonas, for precisely adjusting the antenna composition of photosystem II (Mussgnug et al., 2005; Blifernez-Klassen et al., 2021). Similarly, *LHCBM4* is recognized by NAB1, also presenting a strong RPF peak over the initiation codon, whereas LHCBM1 (not a target of NAB1) has no over-proportional initiation peak (Figure 5C), despite sharing a 75% amino acid sequence identity with LHCBM6. Translational regulation during *LHC* expression is not unique to Chlamydomonas, as *Lhcb4.2*, *Lhcb4.3* and *Lhcb6* are also subject to post-transcriptional regulation in Arabidopsis (Floris et al., 2013). We searched all transcripts that had a gene body coverage of at least 70% (8757 nucleus-encoded, 66 chloroplast-encoded, 6 mitochondria-encoded) for the occurrence of comparable initiation peaks (average coverage of the first seven codons at least four times higher than the average coverage of the remaining CDS). Around 38% of selected nucleus-encoded transcripts had comparable RPF accumulation peaks near the initiation codon, possibly hinting to more co-translational regulation in the cytosol than previously anticipated. In contrast, only 10% (6 transcripts) of the chloroplast-encoded genes showed such peaks (Figure 5D). However, it cannot be excluded that these peaks present artifacts of CHX treatment of the Chlamydomonas cultures during harvest. Upon establishment of the Ribo-seq protocol, several reports demonstrated that pre-incubation of yeast cultures with CHX may create a bias by inducing increased RPF accumulation at initiation codons, which could be the consequence of stalled elongation but unaffected initiation through CHX treatment (Ingolia et al., 2009; Gerashchenko and Gladyshev, 2014; Sharma et al., 2021). However, a recent direct comparison of CHX-treated and untreated conditions demonstrated that CHX causes no initiation-peak bias in human cells, and that even the strong accumulation in yeast might derive from slightly altered protocols between laboratories (Sharma et al., 2021). For Chlamydomonas, CHX biases should result in general initiation peaks, but these peaks are only found in a fraction of transcripts, independent of their translation activity (e.g. *LHCs*). Thus, varying accumulation of RPFs around the initiation codon may be an interesting indicator for altered initiation regulation between transcripts in the cytosol.

We further assayed, if the RPFs of various sizes, ranging from 24 to 34 nt, are randomly distributed across all chloroplast CDS or if a respective RPF size is transcript specific. In general, the dominant RPF size is 28 nt long and the trimodal size distribution appears widely present for translation of most chloroplast transcripts, even if the overrepresentation of the highly translated transcripts (i.e. *rbcL, psbA)* is normalized out (Figure 6A). While present for most chloroplast transcripts, accumulation of the different RPFs sizes varies between transcripts with some over-representation of smaller or larger RPFs, respectively. For example, *psbA, psbH, psaC, psaJ, petD* and *atpH* have an additional strong peak at 24 nt. In contrast, *rpl20* and *psbJ* have a shift towards a dominant population of RPFs with a size >30 nt (Figure 6B-D). Strikingly, the small RPFs within *psbA* and *atpH* show in interesting patter with increased accumulation in the regions of highest RPF coverage. For *pbsA,* the 24 nt RPFs are most prominent in a section between codon 135 and 150, which is just downstream of the transcript encoding the second transmembrane segment (TMS, Figure 6E). It appears as if elongation severely slows down within this region, favoring the extraction of small RPFs, possibly resulting from ribosomes in an “open-ratchet” formation. Alternatively, the smaller reads may result from local binding of a regulatory factor to the transcript, although the small RPFs have similar features as the longer RPFs. Interestingly, we just recently reported that deletion of the One-Helix-Protein 2, OHP2, locally affects *psbA* translation between initiation and the coding sequence of the 3^rd^ TMS. However, ribosome occupancy downstream of the 3^rd^ TMS appeared normal in the mutant. In addition, the absence of OHP2 results in a rapid degradation of nascent PsbA/D1 protein (Wang et al., 2023). Thus, the observed local translation slow-down might be an important stage for PsbA/D1 biogenesis and co-factor integration and OHP2 might be the driver of this regulation. Similarly, RPF accumulation peaks over the CDS of the first TMS of *atpH,* mostly deriving from fragments with a size of 24-25 nt, again suggesting that translation elongation is reduced here (Figure 6E). It will be interesting, to uncover in future experiments if the local translation slowdown is accompanied or caused by binding of regulatory factors.

**Figure 6:**
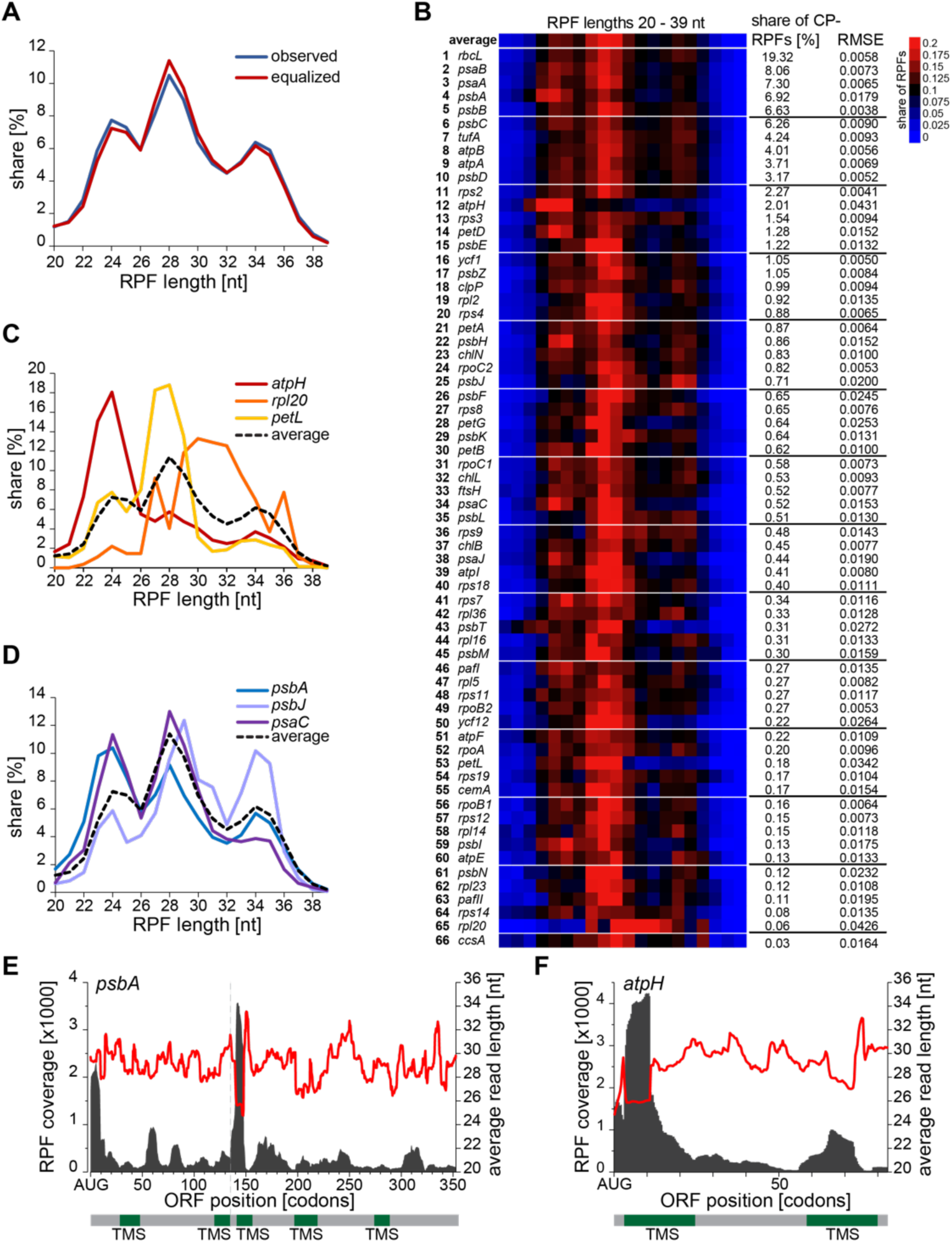
Chloroplast ribosomes generate characteristic ribosome protected fragments. (A) Size distribution of RPFs that map the CDS of chloroplast-encoded transcripts for nuclease treatment conditions iii. (observed). Equalized size distribution shows the normalized size distribution for each chloroplast transcript, neglecting possible weights from the most abundant transcripts. (B) Heatmap presenting the share of the different RPF sizes for each chloroplast transcript. Transcripts are ranked based on their overall share of RPFs relative to all measured chloroplast RPFs. RMSE (Root Mean Square Error) denotes the deviation of the respective size distribution relative to the average size distribution (top row). (C) Size distribution of chloroplast-encoded transcripts with high deviation relative to the average size distribution. (D) Additional size distribution curves of photosynthesis complex subunits. (E) and (F) Ribosome profile of *psbA* and *atpH,* respectively. Red curve shows the RPF sizes, accumulating within the respective CDS section, black curves indicate RFP coverage per nt. Positions of transmembrane segments (TMS) are given below the graph.

### Suitability of ribosome profiles for improving Chlamydomonas genome annotation

Lastly, we aimed to exploit our Ribo-seq data for surveying correct CDS annotations and transcript splice variants within the recently released Chlamydomonas genome, version 6.1 (Craig et al., 2022). In previous genome versions, several genes were falsely annotated, ignoring upstream start codons that were possibly the genuine initiation site of translation (Cross, 2015; Craig et al., 2022). In the new version, the first in-frame ATG codon was generally considered as the initiation codon, in accordance with the scanning model of initiating ribosomes (Hinnebusch, 2011). Systematic comparison of the reference start site with the onset of RPF coverage, confirmed that start codon assignment has been clearly improved between genome version 5.6 and 6.1. However, we detected gene models in which the first ATG appears not to be the genuine initiation site. For example, transcripts encoding LHCA4 (Cre10.g452050) do not initiate from the first ATG (Figure 7A) and were correctly annotated in the previous genome versions. Another example is PRPL3 (Cre10.g417700) (Figure 7B). In addition, we determined some gene models in which RPF accumulation did not correlate with the cognate ATG start codon, possibly hinting to a translation start from non-cognate start sites as exemplified by the *PDI2* CDS (Cre01.g033550, Figure 7C). In this case, the ribosome profile starts with a prominent peak upstream of the annotated start codon located within the annotated 5’-UTR while the annotated start codon is poorly covered compared with the remaining CDS. Here, initiation from the non-cognate TTG could be an explanation (Cao and Slavoff, 2020), which is located in-frame with the annotated coding sequence at two positions within the area covered by the peak. Ribo-seq was already used before to mine for non-cognate start sites in other organisms (Brar and Weissman, 2015). Alternatively, these RPFs could point to the presence of upstream Open Reading Frames for *PDI2*. *PDI2* represents a good example for detecting alternative splicing. Transcript 1 of *PDI2*, possesses a RPF gap between exon 3 and 4 (379 nt downstream of the transcription start site), which is spliced out in transcript versions 2-4 (Figure 7C). In addition, a sudden drop of RPFs is seen at the end of exon 3 in transcript 2 and 4. Based on distribution of RPFs and the transition of ribosome profiles, transcript 5 has the most consistent profile and might hence be the prevalently expressed variant in Chlamydomonas cells, grown under standard laboratory conditions. In future, intelligent modelling and peak deconvolution algorithms could be potentially applied to Ribo-seq data to estimate the stoichiometric ratio of different splice variants’ translation output.

**Figure 7:**
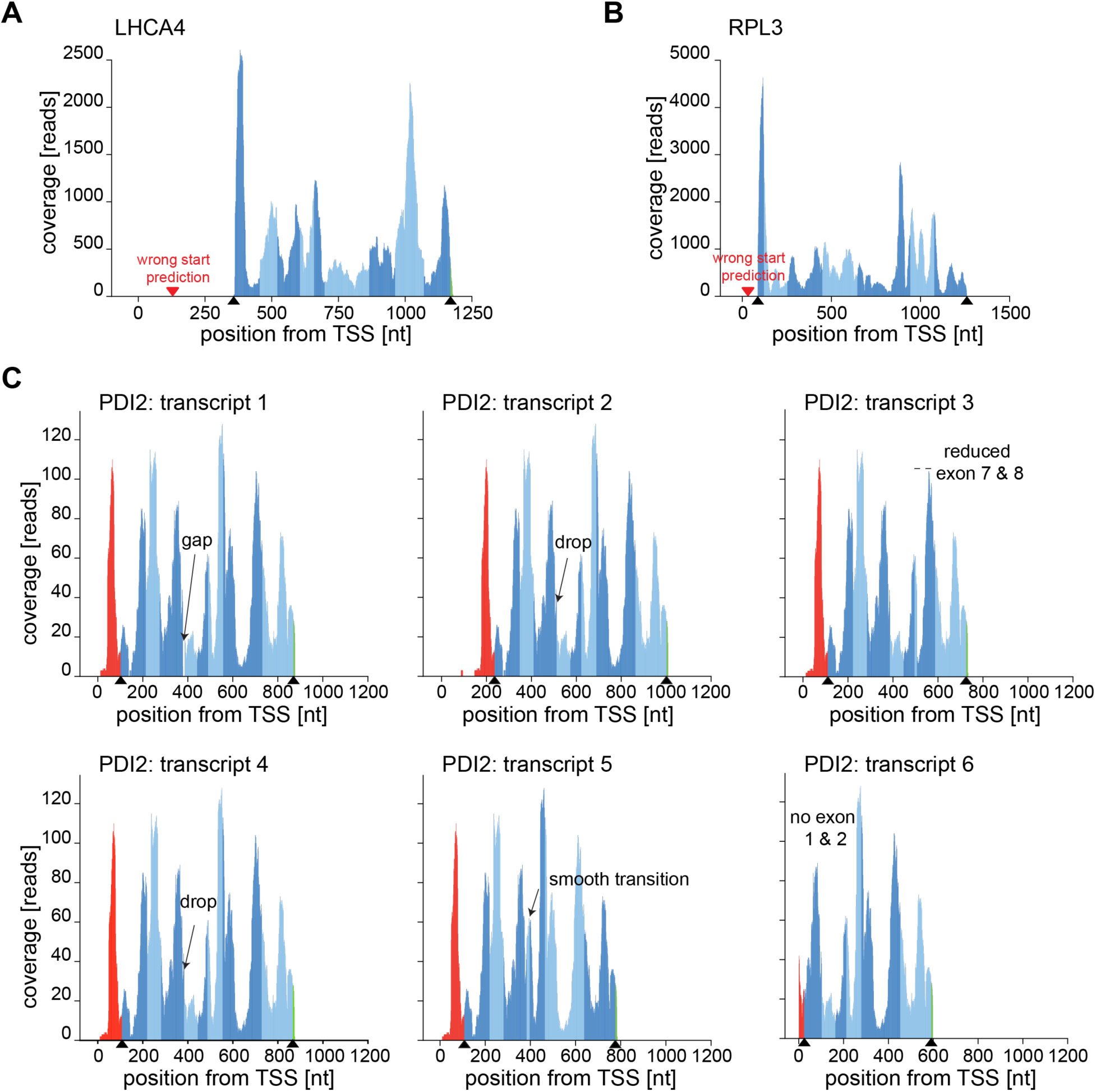
Utilizing ribosome profiles for coding sequence annotation. Exemplary ribosome profiles for determining initiation sites and alternatively spliced transcripts. Red parts of the profiles represent coverage within the 5’-UTRs, green parts represent 3’-UTRs and alternating blue shades represent exons within the CDS. Black arrows on the x-axis indicate start and stop codons. (A) and (B) ribosome profiles of LHCA4 and RPL3 as an example of wrong start codon prediction, respectively. The red arrow indicates the annotated start codon while the black arrows indicate true start and stop codons. (C) Ribosome profiles of PDI2 (Cre01.g033550) transcript variants demonstrating how ribosome profiles can be utilized to identify putative non-cognate start codon initiation and estimate the dominance of transcript variants. Note that AUG was labelled as “true” initiation site, however, an upstream non-cognate start codon might be the correct initiation site.

### Concluding remarks

Taken together, ribosome profiling is a fascinating tool for advancing our understanding of post-transcriptional regulation of gene expression. Our Ribo-seq approach has the resolution and depth to deeply study translation of the nuclear and chloroplast transcripts in Chlamydomonas cells. Furthermore, a highly comparable Ribo-Seq protocol was established for Arabidopsis and tobacco by our team, which will help to directly compare translation between different land plants and Chlamydomonas (Ting et al., 2023). Chlamydomonas is a well-suited organism for understanding system level changes of gene expression throughout the cell cycle or following environmental changes. Ribosome profiles can be of specific help, for a first understanding of the many transcripts, which were not experimentally investigated before. To make our data easily accessible in the research community, we have now linked our Ribo-seq data to the recent Chlamydomonas genome CC-4532 v6.1 (https://phytozome-next.jgi.doe.gov), thereby allowing individuals to conveniently browse the ribosome profiles of their genes of interest. With the new genome release, the number of alternatively spliced transcripts have been significantly expanded. Visualization of ribosome profiles may help to uncover the dominant transcript variants, which are translated under standard laboratory conditions. Certainly, additional experimental approaches are required to understand unexpected features such as translation initiation from non-cognate start sites, RPFs along upstream open reading frames, mechanism of translational regulation, and spatial organization of protein synthesis.

## Materials and Methods

### Cell growth and harvest

*Chlamydomonas reinhardtii* CC-1690 cells were grown mixotrophically in Tris-Acetate Phosphate (TAP) medium (Kropat et al., 2011) to mid-log phase (4 to 5×10^6^ cells per mL) under constant light of 80 mmol m^-2^ s^-1^ (MASTER LEDtube HF 1,200 mm UO 16W830 T8 and 16W840 T8, Philips) and 25°C on a rotary shaker. Immediately before harvest, 100 µg/mL chloramphenicol (CAP) and cycloheximide (CHX) was added and cells were rapidly chilled by pouring over -80°C cold silicon ice cubes until 4°C culture temperature was reached. Subsequently, cells were pelleted by 2 min centrifugation at 4,000 g and 4°C, washed in ice-cold TKM+ buffer (50 mM acetate buffered Tris pH 8.0, 150 mM KCl, 10 mM MgCl_2_, 100 µg/mL CAP, 100 µg/mL CHX) and pelleted again using the same centrifugation conditions. Afterwards, the pellet was resuspended in freezing buffer (TKM+ supplemented with 100 mM phenyl methyl sulfonyl fluoride (PMSF) and 16% of 25x Roche protease inhibitor cocktail) by slow pipetting in 1/2000 of the initial culture volume and flash frozen by dripping the cell suspension into liquid nitrogen. Frozen cells were stored at -80°C until further use.

### Ribosome profiling and RNA sampling

Frozen cells were ground using a bead mill with nitrogen-cooled steel containers and beads for two times 2 min at 27 Hz frequency and cooling in liquid nitrogen between rounds. The cell powder was mixed with equal volumes of 2x concentrated lysis buffer (2x concentrated TKM+ buffer supplemented with 2 mM dithiotreitol (DTT), 2% Triton X-100 and 20% sucrose) and incubated for 5 min at 4°C before pelleting cell debris at 8,000 g, 4°C for 10 min. For nucleolytic digest conditions i. and ii., lysate was layered on a 2.5 mL 64% sucrose cushion prepared in TKM+ buffer supplemented with 1 mM DTT and ultracentrifuged for 3 h at 60,000 rpm and 4°C in a Ti70 rotor. The resulting polysome pellet was briefly rinsed with 500 µL ice-cold diethyl pyro carbonate (DEPC)-treated water and resuspended overnight in 100 µL ice-cold ribosome buffer (2x concentrated TKM+ buffer diluted with 25x concentrated Roche protease inhibitor cocktail to a total amount of 16%, 1 mM PMSF and 0.1% Triton X-100). Remaining insoluble debris were removed by centrifugation for 1 min at 1,500 g before addition of 1 U of RNase I (Ambion) per µg RNA and 0.134 U TURBO DNase (Invitrogen). Nucleolytic digest was performed for 1 h at 4°C (condition i.) or 23°C (condition ii.) and stopped by addition of 0.4 U per applied unit of RNase I of SUPERase•In RNase Inhibitor (Invitrogen). For condition iii., 1 U of Ambion RNase I (Invitrogen) per µg RNA and 0.134 µL per µL lysate of TURBO DNase (Invitrogen) was directly added to the lysate and incubated for 1 h at 4°C. The digest was stopped by addition of 0.4 U of SUPERase•In RNase Inhibitor (Invitrogen) per applied unit of RNase I. The lysate was centrifuged again for 10 min at 8,000 g and 4°C to remove any cell debris that may have precipitated during the nuclease digest. For all conditions, monosomes were pelleted through a 750 µL 30% sucrose cushion prepared in TKM+ buffer supplemented with 1 mM DTT for 30 min in a S150AT rotor at 72,000 rpm and 4°C. The resulting ribosome pellets were resuspended in 100 µL of ice-cold ribosome buffer, then supplemented with 15 mM ethylene diamine tetra-acetic acid (EDTA), pH 8.0 and instantly mixed with 750 µL Invitrogen TRIzol reagent (Invitrogen) and incubated at room temperature for 5 min. The solution was mixed with 150 µL chloroform and incubated for 2 min at room temperature before 15 min centrifugation at 20,000 g and 4°C. The supernatant was mixed with 3 µL GlycoBlue co-precipitant (Invitrogen), 1:10 volume of 3 M sodium acetate, pH 5.5, 1 volume isopropanol and incubated overnight at -20°C. The precipitated RNA was pelleted by 1 h centrifugation at 20,000 g and 4°C, briefly washed with ice-cold 80% ethanol and centrifuged again for 30 min at the same parameters. The resulting RNA pellet was briefly dried and dissolved in 11 µL of nuclease-free water.

For RNA-seq samples, 10 mL of culture were harvested by 2 min centrifugation at 4,000 g at room temperature (before harvesting the culture for RPF isolation), shock-freezing in liquid nitrogen and storage at -80°C. Samples were lysed in prewarmed (50°C) lysis buffer (50 mM Tris-HCl pH 8.0, 200 mM NaCl, 20 mM EDTA pH 8.0 and 2% SDS) and incubated for 2 min at 50°C. Immediately after incubation, 500 µL of TRIzol (Invitrogen) reagent were added and the mixture was incubated for 5 min at room temperature. The suspension was mixed with 200 µL chloroform and incubated for another 5 min at room temperature before 3 min centrifugation at 12,000 g and room temperature. The nucleic acid containing phase was transferred to a fresh tube and mixed with 1.5 volumes of ethanol. RNA was purified by the Macherey Nagel NucleoSpin RNA Plant kit.

### Ribosome protected fragment extraction and rRNA depletion

50 µg RNA from the ribosomal extractions were diluted to a volume of 20 µL with nuclease-free water, mixed with 2x formamide RNA loading buffer (90% deionized formamide, 20 mM Tris-HCl, pH 7.5, 0.5 M EDTA, 0.04% bromophenol blue in ethanol) and denatured for 90 s at 80°C. The samples were instantly put on ice and then loaded onto 12% urea-TBE-gels (90 mM Tris, 9 mM boric acid, 2 mM EDTA, 12% 19:1 40% acrylamide-bisacrylamide mix, 8 M urea). Gels were run at 200 V for approximately 1 h and incubated with SYBR Gold nucleic acid stain (Invitrogen), diluted 1:10,000 in TBE buffer (90 mM Tris, 9 mM boric acid, 2 mM EDTA). RPF were excised from the gels in a size range from 20 – 40 nt according to a RNA size marker. Gel pieces were crushed and incubated overnight with 400 µL RPF elution buffer (300 mM sodium acetate, pH 5.5, 0.25% sodium dodecyl sulfate and 1 mM EDTA, pH 8.0) at room temperature. The eluate was collected, mixed with 3 µL GlycoBlue co-precipitant, 1 volume of isopropanol and incubated overnight at -20°C. The precipitated RNA was pelleted by 1 h centrifugation at 20,000 g and 4°C, briefly washed with ice-cold 70% ethanol and centrifuged again for 30 min at the same parameters. The resulting RNA pellet was briefly dried and dissolved in 20 µL of nuclease-free water. For Ribo-seq samples, abundant rRNA fragments were depleted by incubating 100 - 200 ng of ribosome RPFs with 1.19 µL of 100 µM biotinylated rRNA-oligo depletion mix (consisting of the oligonucleotides: AATATGCGTTCAAAGATTCGATGATTCACG, TAGCTCTAGAATTACTACGGTTATCCGAGTA, TACCCGACGCTGAGGCAGACATGCTCTTGG, GATTCGTGAAGTTATCATGATTCACCGCA, ACGGGATGAATCTCAGTGGATCGTAGCA, CGATCTAGCCGTCTTAGAGCTAGAAGCAGG) with 4 µL formamide, 1 µL 20x concentrated hybridization buffer (300 mM sodium citrate, 3 M NaCl), 2 µL 0.5 M EDTA, pH 8.0 and in a total volume of 20 µL. The mixture was hybridized in a thermocycler by heating up to 80°C for 5 min and subsequent cooling in 5°C steps for 2 min each until 35°C was reached in the device. Afterwards, 60 µL of hybridization buffer, supplemented with 20% formamide was added to the tube and the sample was depleted twice for 15 min using 15 µL of Dynabeads MyOne Streptavidin T1 magnetic beads (Invitrogen) freshly washed and prepared according to the manufacturer’s manual. The supernatant was mixed with 3 µL GlycoBlue co-precipitant, 2.5 volumes of ethanol and incubated overnight at -20°C. RPFs were pelleted by 1 h centrifugation at 20,000 g and 4°C, briefly washed with ice-cold 70% ethanol and centrifuged again for 10 min at the same parameters. The resulting pellet was briefly dried and dissolved in 41 µL of nuclease-free water. Excess oligos were removed by 30 min DNase digest at 37°C upon addition of 5 µL 10x TURBO DNase buffer, 2 µL Invitrogen TURBO DNase and 2 µL SUPERase•In RNase Inhibitor (Invitrogen). After digest, RPFs were mixed with 150 µL of 100 mM NaCl and purified again by phenol-chloroform precipitation and final precipitation overnight at -20°C using 2.5 volumes of ethanol and 2 µL GlycoBlue co-precipitant. RPFs were pelleted by 1 h centrifugation at 20,000 g and 4°C, briefly washed with ice-cold 70% ethanol and centrifuged again for 10 min at the same parameters. The resulting pellet was briefly dried and dissolved in 36.5 µL of nuclease-free water.

For sequencing library preparation, ribosome RPFs were denatured at 65°C for 5 min in a thermocycler and immediately transferred to ice. The sample was then mixed with 5 µL 10 x T4-polynucleotide kinase buffer A, 2.5 µL T4-polynucleotide kinase (Thermo Scientific), 2 µL SUPERase•In RNase Inhibitor and incubated for 10 min at 37°C. Subsequently, 5 µL of 10 mM ATP was added and the sample was incubated for additional 30 min at 37°C. After phosphorylation, the RPFs were mixed with 150 µL 100 mM NaCl and purified again by phenol-chloroform precipitation and final precipitation overnight at - 20°C using 2.5 volumes of ethanol and 2 µL GlycoBlue co-precipitant. RPFs were pelleted by 1 h centrifugation at 20,000 g and 4°C, briefly washed with ice-cold 70% ethanol and centrifuged again for 10 min at the same parameters. The resulting pellet was briefly dried and dissolved in 11.5 µL of nuclease-free water. The whole sample was used as input for the NEXTFLEX Small RNA-seq Kit v3 (Perkin Elmer) to generate libraries according to the kit’s manual. For RNA-seq, the purified RNA was later used as input for the Zymo-Seq RiboFree Total RNA library kit for library preparation according to the manufacturer’s manual.

### Processing of raw sequencing data

Demultiplexed FASTQ files were derived from an Illumina NextSeq 550 system aiming for 40 M reads per Ribo-seq sample and 20 M reads per RNA-seq sample at a read length of 75 nt. Samples were processed through a custom-made pipeline in form of a Linux shell script calling several bioinformatic tools and custom Python scripts. First operation of the pipeline was removing 3’ adapter sequences using cutadapt for RNA-seq and Ribo-seq, respectively (Martin, 2011) with parameters [--minimum-length=9, - a=”TGGAATTCTCGGGTGCCAAGG”]. After removal of 3’ adapter sequences, the 3’ and 5’ unique molecular identifier tags (UMI tags) were removed from each read’s sequence and shifted to the rear of the header row by UMI tools (Smith et al., 2017) with parameters [extract --bc-pattern=NNNN] for 5’ UMI tags and [extract --bc-pattern=NNNN –3prime] for 3’ UMI tags. Remaining reads were filtered for sequence length using cutadapt again with parameters [--minimum length=20 --maximum-length=39] to remove any read that is unlikely to represent a real RPF. After size filtering, a custom-made Python script corrected the UMI-tags that were previously shifted to the read headers in the form [header line_NNNN_NNNN] to [header line_NNNNNNNN], allowing UMI tools in a later step to deduplicate the data set based on both UMI tags. In a first mapping step, reads were aligned to a set of *C. reinhardtii* non-coding RNA FASTA sequences provided by ENSEMBL plants together with the genome version 5.5 to remove contaminating RNA species using STAR with parameters [--runMode alignReads --outSAMtype BAM SortedByCoordinate --outFilterMultimapNmax 20 --outFilterMismatchNmax 3 -- outReadsUnmapped Fastx --alignIntronMax 7000 --twopassMode Basic]. Reads that did not align to the ncRNA set were copied to a new FASTQ file that was mapped against the *C. reinhardtii* genome version 6.1 using STAR (Dobin et al., 2013) with parameters [-- runMode alignReads –outSAMtype BAM SortedByCoordinate --outFilterMismatchNoverLmax 0.1 --outReadsUnmapped Fastx -- alignIntronMax 8000 –outSAMmultNmax 1 –outMultimapperOrder Random]. After genome mapping, the resulting BAM file was deduplicated using UMI-tools’ dedup function and afterwards indexed using samtools’ index function (Li et al., 2009).

### Data processing

All analyses on the NGS data sets including calculations of gene body coverage, periodicity, P-site offsets, organellar read length distributions, ribosome profiles and gene expression metrics were performed using a custom-made object-oriented Python module that makes extensive use of the packages pysam/HTSlib (https://github.com/pysam-developers/pysam), pandas (McKinney, 2010), NumPy (Harris et al., 2020), SciPy (Virtanen et al., 2020), matplotlib (Hunter, 2007) and seaborn (Waskom, 2021) among multiple others. For RPKM calculations in Ribo-seq data sets, only reads mapping to the annotated CDS of a transcript were considered. Alternatively we calculate counts per million values for the interested reader (Supplemental Dataset). For any calculations on RNA-seq data sets, reads mapping to the whole transcript were considered and the full length of the mature transcript was used for RPKM calculations. Translation efficiency values were simply calculated by dividing an CDSs RPKM value from a Ribo-seq data set by the respective transcripts RPKM value from the affiliated RNA-seq data set.

For the 5’-P-site / 3’-A-site offset calculations, all reads fully enclosing either an annotated start codon (5’-P-site offset) or an annotated stop codon (for 3’-A-site offset) were taken into consideration separately for the nuclear and the chloroplast genome. These reads were sorted according to the length of their alignment to the genome, which was considered as read length to avoid miscalculations due to clipped or masked sequences. For every read length species, the position of the reads 5’ end or 3’ end relative to the position of the first base of the start codon (for 5’-P-site offsets) or the last base of the stop codon (for 3’-A-site offsets) was determined as the offset. Afterwards, the most frequently occurring offset for a read length species was determined to be the true offset of this species and was used in the P-site calculation. To display the results of the offset calculations, the count matrix of both 5’-P-site offsets and 3’-A-site offsets were normalized separately to the total number of counts each and plotted as heatmap.

For calculation of the P-sites first position of reads mapping to nucleus-encoded CDSs, all reads mapping to a CDS were filtered from the data set and for every read, the 5’-most mapping position of the read’s alignment + the 5’-P-site offset determined for the reads alignment length was determined to be the first P-site position in case of transcripts encoded on the + strand. For reads encoded on the minus strand, the first P-site position was determined to be the 3’-most mapping position of the RPF alignment minus the determined 5’-P-site offset. In all cases, splicing was considered and whenever a spliced read was found, the calculation was adjusted according to the number of genomic positions spanned by the splice junction of the read. For reads mapping to chloroplast-encoded transcripts on the + strand, the first P-site position was determined to be the 3’-most mapping position – (3’-A-site offset + 5). For reads mapping to chloroplast-encoded transcripts on the - strand, the first P-site position was calculated to be the 5’-most mapping position + (3’-A-site offset + 5). The metagene P-site profiles of nucleus-encoded transcripts were calculated considering only the 2,000 most expressed transcripts of a dataset, according to their RPKM values. If multiple splice variants of the same gene occurred in the list, only the longest variant (annotated as transcript 1 in the genome) was considered for the calculation to avoid analyzing the same reads multiple times. The profiles were constructed by calculating the first position of the P-site for all reads mapping to the first 30 and the last 30 codons of considered transcript CDSs. For a ribosome RPF to be in-frame with the annotated CDS, the first position of the P-site must be located on the first base of a codon. According to this logic, the calculated P-site positions were categorized into “in-frame”, “+1 shifted” and “+2 shifted” and summed up for all codons of considered transcript CDSs. To calculate the frame preference per codon, the counts in each category in a codon were divided by the sum of all categories for every codon separately. For calculation of the chloroplast metagene P-site profile, all protein-coding chloroplast transcripts were considered.

The extension of gene body coverage versus a respective cut-off was calculated as the portion of an CDS that was covered by the number of reads equal to or greater than the respective cut-off. This procedure was applied to all annotated CDSs using increasing cut-offs from 1 to 10 reads to monitor the stepwise decrease of gene-body coverage. For every applied cut-off, the CDSs were categorized as being covered over >10% of their length to >90% of their length in 10% steps and CDSs falling into every category were counted and displayed in a heat map to estimate both completeness and strength of CDS coverage in the data set.

To calculate the read length distribution, all reads mapping to the respective genome were filtered and the length of their alignment to the genome was taken as read length. After counting the number of all possible read lengths for each genome separately, these numbers were normalized by the total number of reads mapping to the respective genome. For read biotype distribution, the reads were further categorized if they map to an annotated coding sequence, 5’- or 3’-UTR or intergenic region. To handle reads overlapping two regions of different category, for example 5’-UTR and coding sequence as seen regularly in initiation peaks, the read was assigned the category that had the larger share on the read’s alignment. In rare cases, when a read mapped to different categories in two different splice variants of the same transcript, the read was assigned to both categories.

## Supplemental data

**Supplemental Figure S1**: Features of ribosome protected fragment coverage.

**Supplemental Figure S2:** RPF length distribution and periodicity.

**Supplemental Figure S3:** Transcript accumulation, translation output and translation efficiency of nuclear genes.

**Supplemental Figure S4:** Correlation between nuclease digestion conditions, transcript accumulation and translation.

**Supplemental Figure S5:** Correlation analysis of RPF count between nuclease digest conditions.

**Supplemental Figure S6:** Detection of abnormal initiation peaks in the three genomes of *Chlamydomonas reinhardtii*.

**Supplemental Dataset.** List of accession numbers, gene body coverage, RPKM and CPM of Ribo-seq and RNA-seq experiments.

## Acknowledgements

We thank Fabian Ries, Claudia Herkt and Peter Emelin for critical discussion of the data and the manuscript. Martin Simon, Jaro Schmitt and Günter Kramer for help with next generation sequencing. We further thank Jeremy Phillips and David Goodstein from the Joint Genome Institute for making our data available for the public on the Phytozome website.

## Author contribution

V.L.G. performed ribosome profiling experiments, bioinformatic data analyses and wrote parts of the manuscript. M.K.Y.T. and R.Z. helped with setting up ribosome profiling, N.H. and S.R. conducted correlation analyses. F.W. designed experiments, analyzed data, and wrote the manuscript.

## Funding

This work was supported by the Deutsche Forschungsgemeinschaft grant 437345987 (WI 3477/3-1, ZO 302/5-1, RU 2266/2-1 to F.W., R.Z. and S.R., respectively) and the Forschungsschwerpunkt BioComp to F.W.

## Data availability

All gene accession codes are given in the Supplemental Dataset and are in accordance with the latest Chlamydomonas genome v6.1 annotation (https://phytozome-next.jgi.doe.gov) or the GenBank/EMBL data libraries. The full Ribo-seq and RNA-seq data are accessible via the Dataplant consortium.

## Competing interests

The authors declare no competing interests.

## SUPPLEMENTAL INFORMATION

### Supplemental Figures and Legends

**Supplemental Figure S1:**
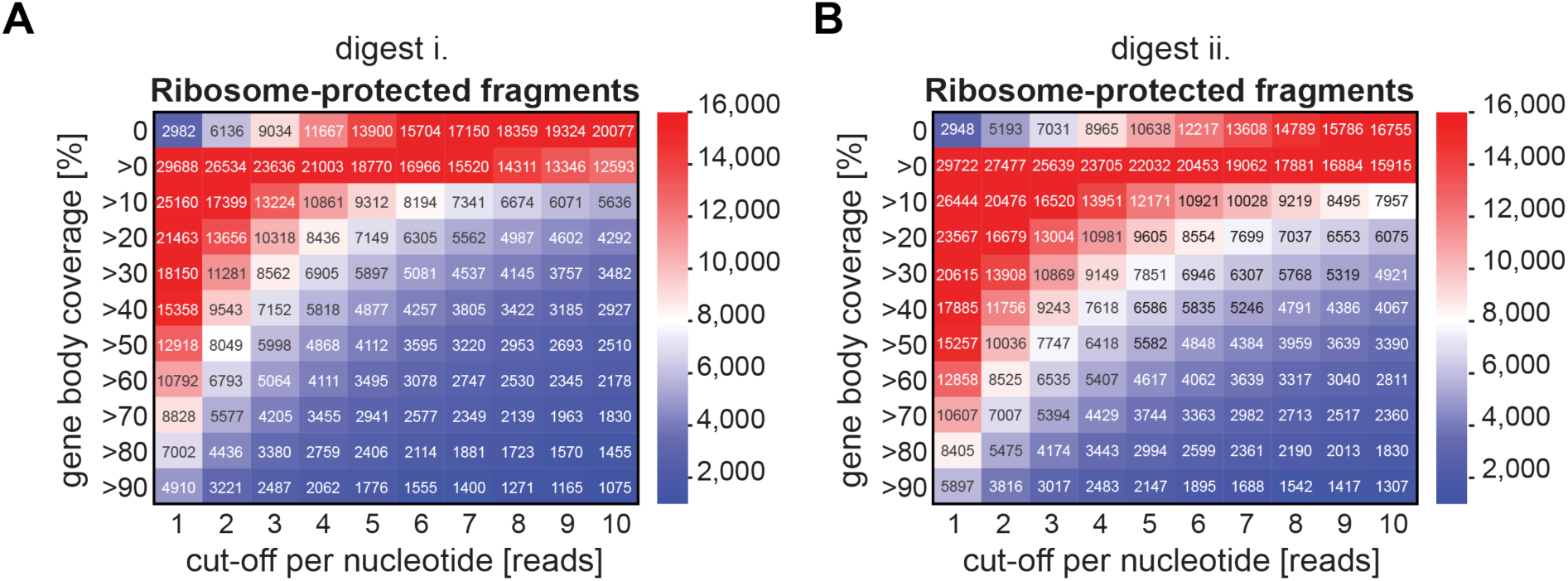
Features of ribosome protected fragment coverage. Supporting Figure 1. (A) and (B) heatmaps representing transcript numbers categorized by their extent of gene body coverage in dependence of specific read count cut-offs for Ribo-seq data sets i. and ii.

**Supplemental Figure S2:**
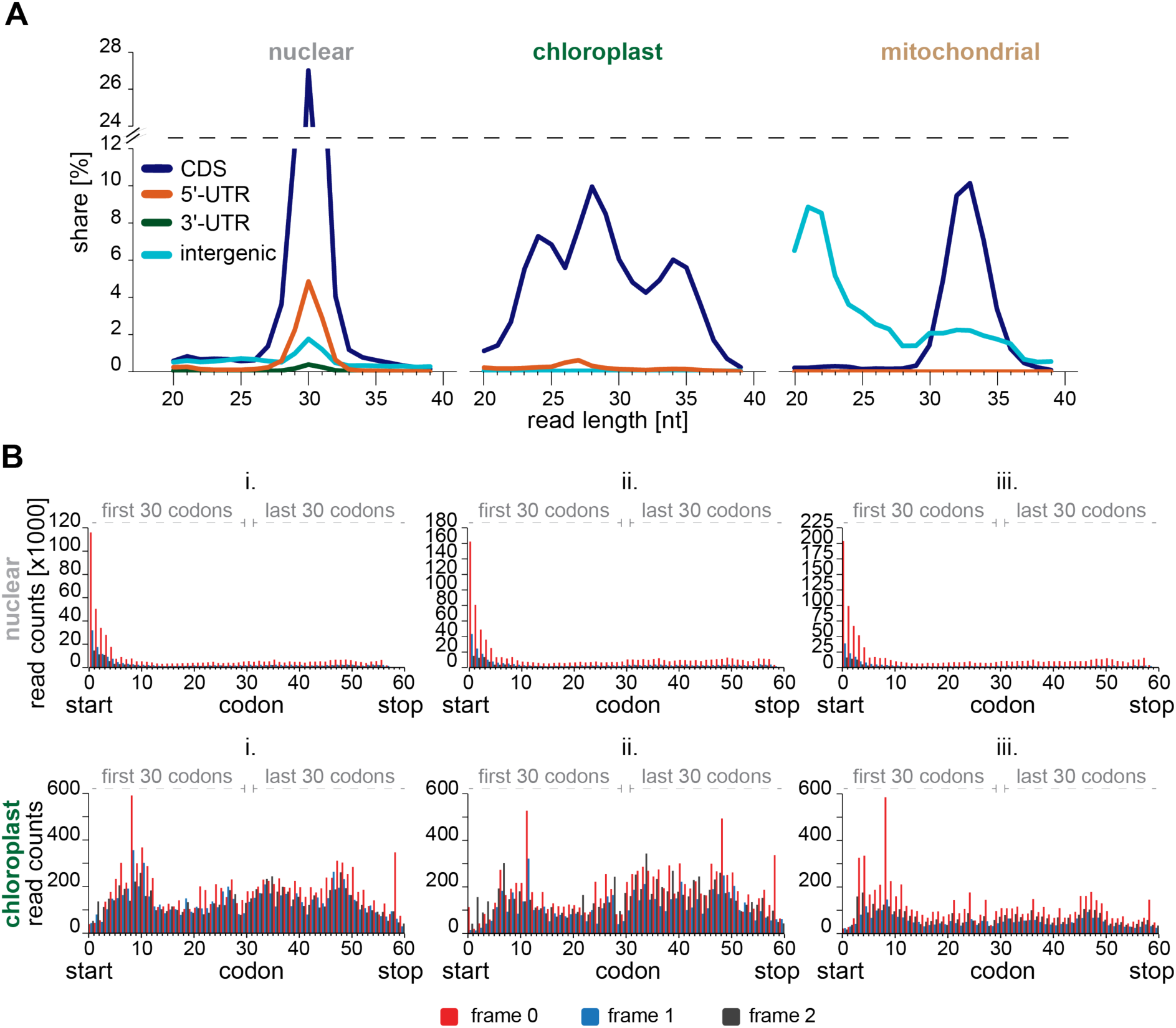
RPF length distribution and periodicity. Supporting Figure 2. (A) Length distribution of RPFs for each organelle separated by their match to CDS, 3’- or 5’-UTR or intergenic regions (only data of digest iii. are shown here). (B) metagene RPF accumulation of nuclear and chloroplast-encoded transcripts per codon for all three nuclease treatments. Codons 1-30 represent the first 30 codons of each transcript, whereas codons 31-60 represent the last 30 codons of each transcript. Frame 0 corresponds to AUG in the ribosomal P-site.

**Supplemental Figure S3:**
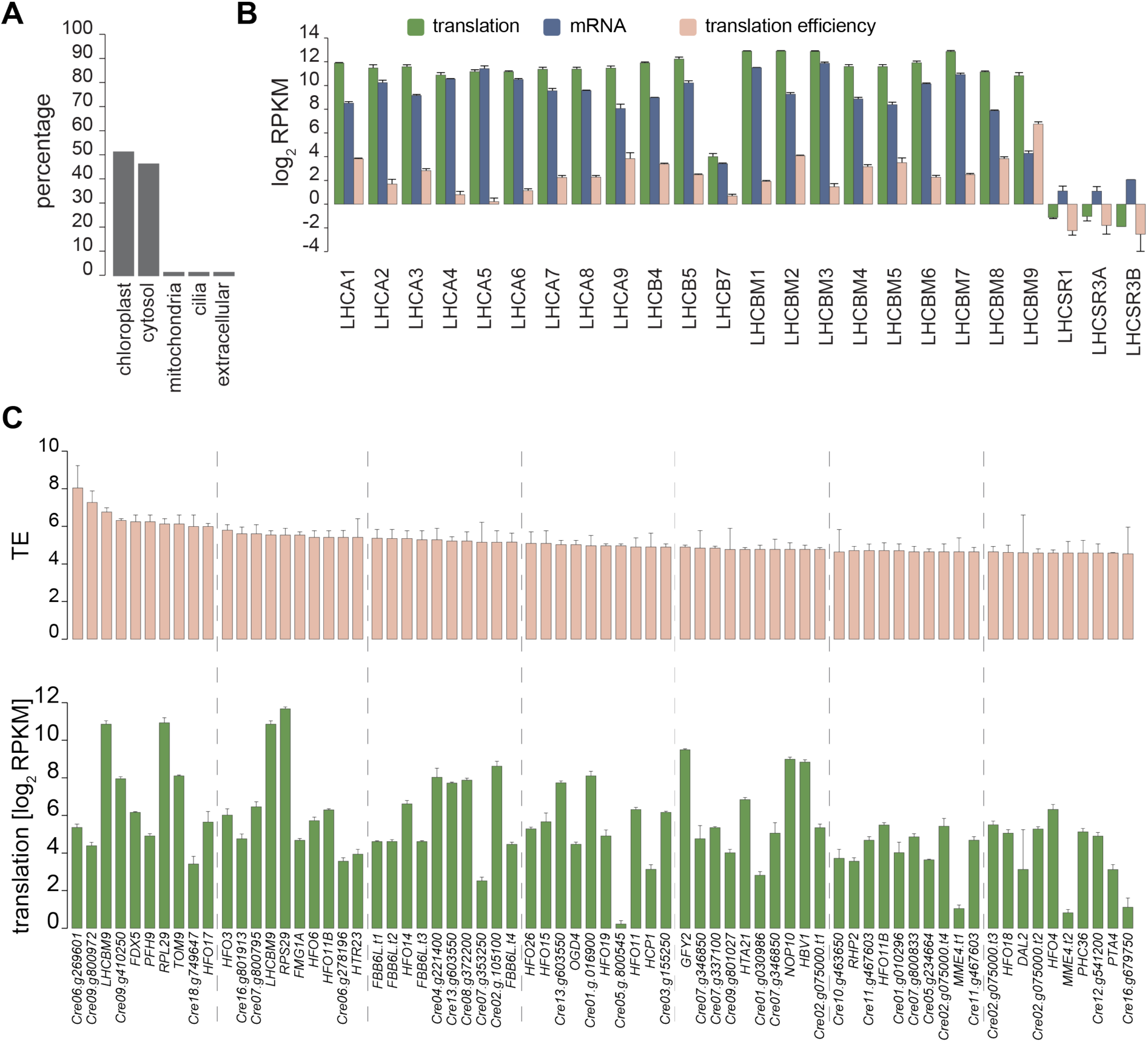
Transcript accumulation, translation output and translation efficiency of nuclear genes. Supporting Figure 4. (A) Subcellular distribution of the top 100 expressed nucleus-encoded transcripts on the translation level. If multiple transcript variants of the same gene occurred in the list, only the longest occurring variant was considered. (B) Translation output, transcript abundance and translation efficiency of light harvesting protein encoding transcripts across all three Ribo-seq data sets, considering only the longest transcript variant, respectively. Mean values averaging all three Ribo-seq data are plotted, error bars denote standard deviations. (C) Translation efficiency and translation output of the 70 most efficiently translated nucleus-encoded transcripts. Mean values averaging all three Ribo-seq data are plotted, error bars denote standard deviations. If multiple transcript variants of the same gene occurred in the list, only the longest occurring variant was considered, unless stated in the legend.

**Supplemental Figure S4:**
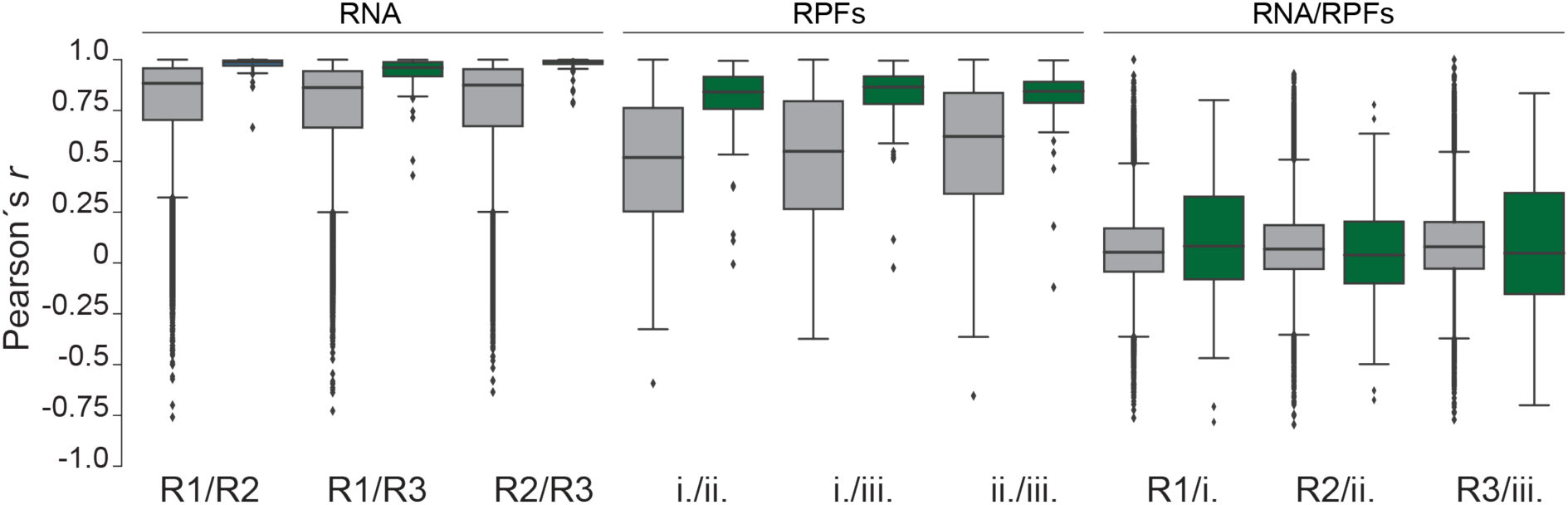
Correlation between nuclease digestion conditions, transcript accumulation and translation. Supporting Figure 5. Boxplots representing the distribution of Pearson correlation coefficients between ribosome profiles and RNA-seq read distribution of the same transcripts each in different data sets. Boxes represent the 0.25 to 0.5 and the 0.5 to 0.75 quartiles, lines represent the median values and whiskers represent the remaining data. Outliers are defined as values being smaller or greater than the respective median ±1.5 times the interquartile range and are depicted as points R1/2/3 indicate different RNA-seq data sets.

**Supplemental Figure S5:**
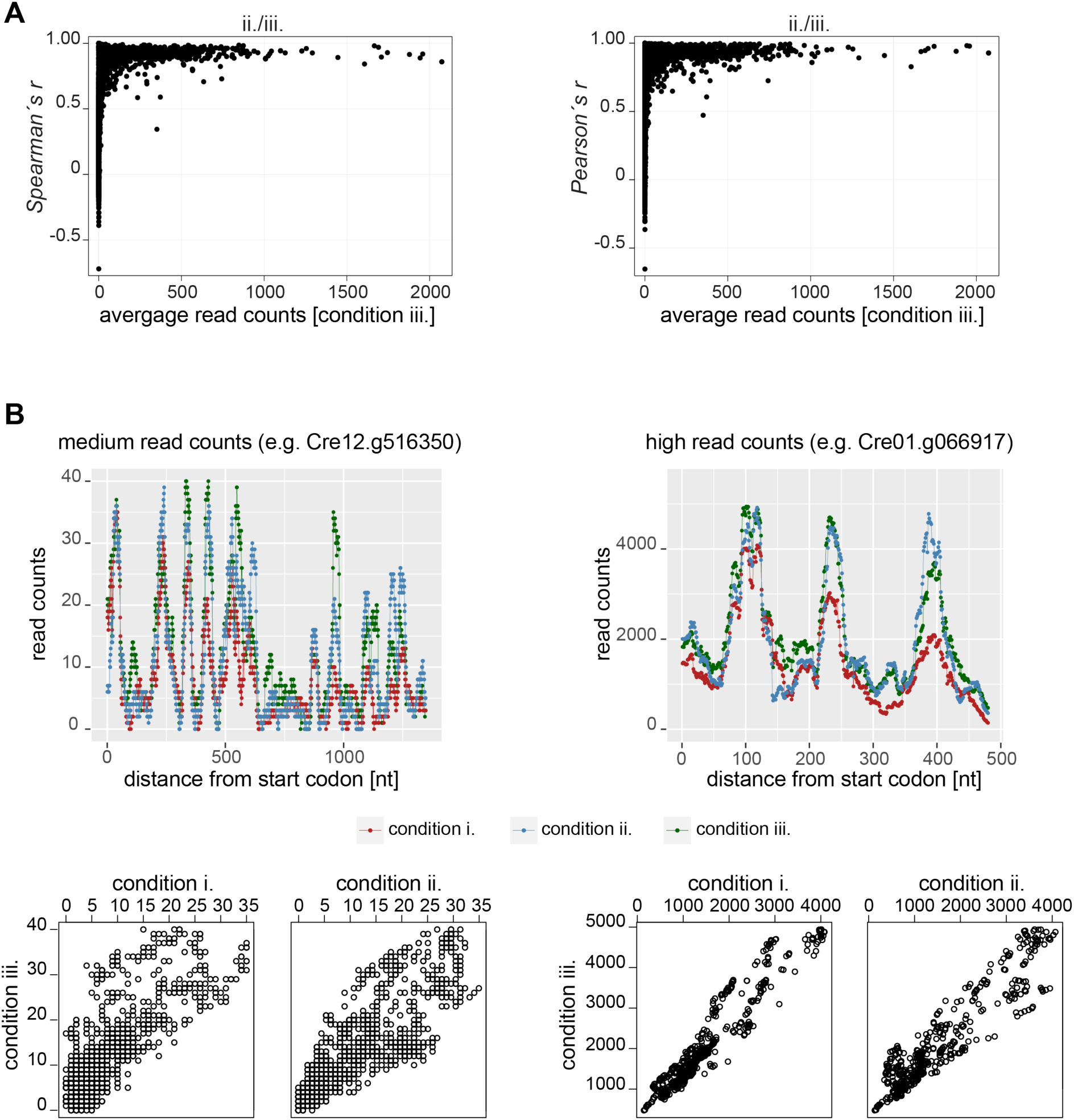
Correlation analysis of RPF count between nuclease digest conditions. Supporting Figure 5. (A) Relation between correlation coefficient comparing two experimental conditions and average read counts. For each transcript, counts were summed over all positions and then divided by the total number of nucleotides in the transcript. The correlation is high unless the average read counts are low. The distribution of *r*-values might be well suited for selecting good cut-off ranges for a deeper analyses of ribosome profiles. (B) Examples of ribosome profiles with medium and high read counts for all three experimental conditions. Scatter plot of read counts for each nucleotide for comparison of two experimental conditions. Perfectly identical profiles would correspond to a straight line in the scatter plot.

**Supplemental Figure S6:**
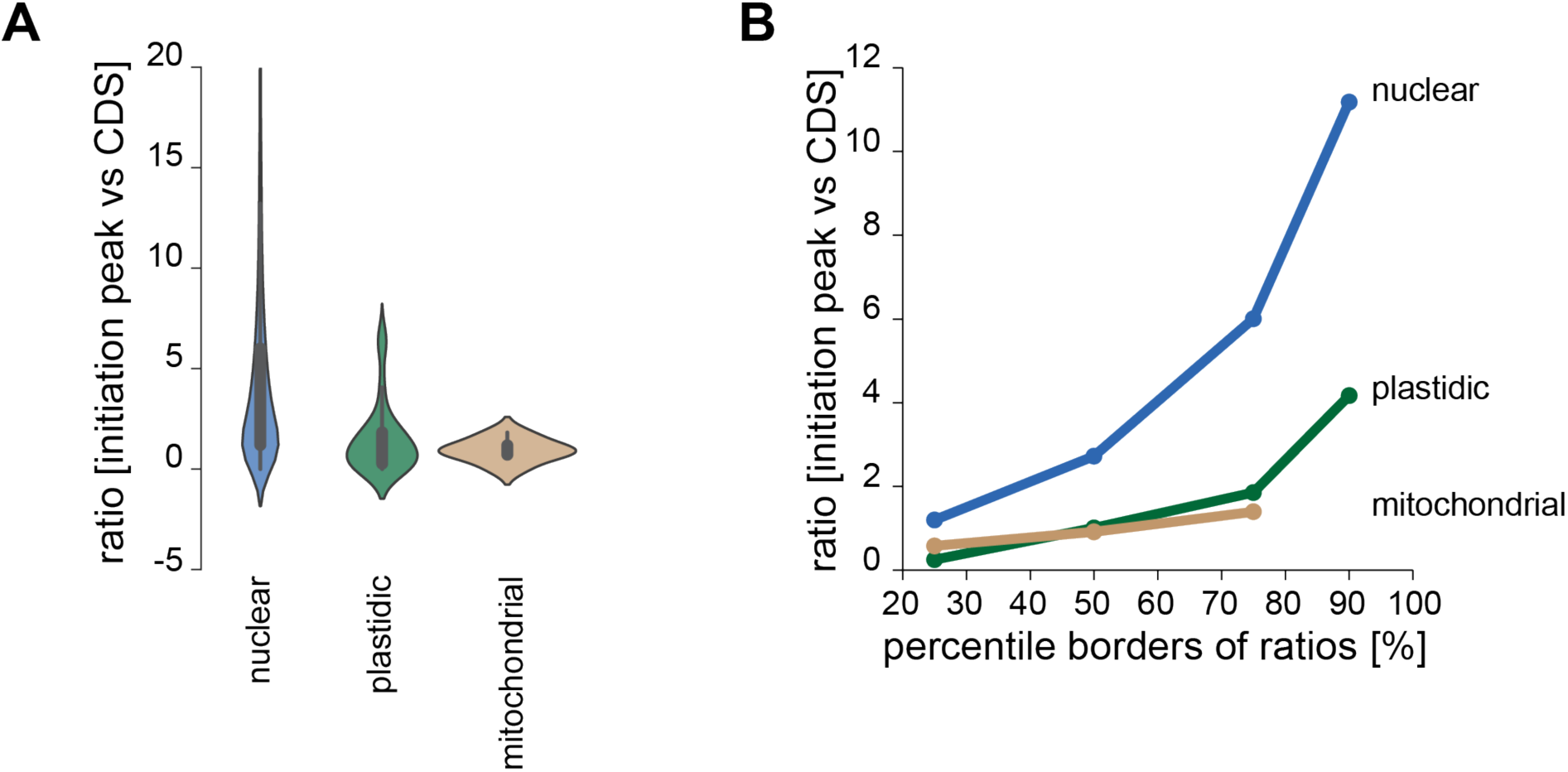
Detection of abnormal initiation peaks in the three genomes of *Chlamydomonas reinhardtii*. Supporting Figure 5. (A) Violin plots representing the distribution of ratios between average per nucleotide coverage within the first seven codons and the average per nucleotide coverage of the remaining CDS for each organelle’s transcripts separately. Thick lines within the violins represent the interquartile range while thin lines represent the whiskers with a cut-off of 1.5 times the interquartile range. (B) Scatter plot with lines representing the increase of the percentile borders of the distributions shown in (A). In both cases, only transcripts with a gene-body coverage of at least 70% were considered (8757 nucleus-encoded, 66 chloroplast-encoded, 6 mitochondrial-encoded) to minimize the risk of misinterpretations due to low or absent coverage in either of both regions. Calculation of a 90^th^-percentile border for mitochondrial transcripts was not possible due to the low number of transcripts in consideration.

## Notes

### Competing Interest Statement

The authors have declared no competing interest.

### Summary of Updates

We updated the calculation of the translation output (RPKM) and we added a new Figure 6. The manuscript has now 7 main figures.

## References

Aird D, Ross MG, Chen WS, Danielsson M, Fennell T, Russ C, Jaffe DB, Nusbaum C, Gnirke A (2011) Analyzing and minimizing PCR amplification bias in Illumina sequencing libraries. Genome Biol 12: R18

Blifernez-Klassen O, Berger H, Mittmann BGK, Klassen V, Schelletter L, Buchholz T, Baier T, Soleimani M, Wobbe L, Kruse O (2021) A gene regulatory network for antenna size control in carbon dioxide-deprived *Chlamydomonas reinhardtii* cells. Plant Cell 33: 1303–1318

Bonente G, Pippa S, Castellano S, Bassi R, Ballottari M (2012) Acclimation of *Chlamydomonas reinhardtii* to different growth irradiances. J Biol Chem 287: 5833–5847

Brar GA, Weissman JS (2015) Ribosome profiling reveals the what, when, where and how of protein synthesis. Nat Rev Mol Cell Biol 16: 651–664

Calviello L, Mukherjee N, Wyler E, Zauber H, Hirsekorn A, Selbach M, Landthaler M, Obermayer B, Ohler U (2016) Detecting actively translated open reading frames in ribosome profiling data. Nat Methods 13: 165–170

Cao X, Slavoff SA (2020) Non-AUG start codons: Expanding and regulating the small and alternative ORFeome. Exp Cell Res 391: 111973

Chartron JW, Hunt KC, Frydman J (2016) Cotranslational signal-independent SRP preloading during membrane targeting. Nature 536: 224–228

Chen H, Alonso JM, Stepanova AN (2022) A Ribo-Seq Method to Study Genome-Wide Translational Regulation in Plants. Methods Mol Biol 2494: 61–98

Chiu CW, Li YR, Lin CY, Yeh HH, Liu MJ (2022) Translation initiation landscape profiling reveals hidden open-reading frames required for the pathogenesis of tomato yellow leaf curl Thailand virus. Plant Cell 34: 1804–1821

Chotewutmontri P, Barkan A (2016) Dynamics of chloroplast translation during chloroplast differentiation in maize. PLoS Genet 12: e1006106

Chotewutmontri P, Barkan A (2018) Multilevel effects of light on ribosome dynamics in chloroplasts program genome-wide and *psbA*-specific changes in translation. PLoS Genet 14: e1007555

Chotewutmontri P, Barkan A (2020) Light-induced *psbA* translation in plants is triggered by photosystem II damage via an assembly-linked autoregulatory circuit. Proc Natl Acad Sci U S A 117: 21775–21784

Chung BY, Hardcastle TJ, Jones JD, Irigoyen N, Firth AE, Baulcombe DC, Brierley I (2015) The use of duplex-specific nuclease in ribosome profiling and a user-friendly software package for Ribo-seq data analysis. RNA 21: 1731–1745

Conesa A, Madrigal P, Tarazona S, Gomez-Cabrero D, Cervera A, McPherson A, Szczesniak MW, Gaffney DJ, Elo LL, Zhang X, Mortazavi A (2016) A survey of best practices for RNA-seq data analysis. Genome Biol 17: 13

Costanzo MC, Hogan JD, Cusick ME, Davis BP, Fancher AM, Hodges PE, Kondu P, Lengieza C, Lew-Smith JE, Lingner C, Roberg-Perez KJ, Tillberg M, Brooks JE, Garrels JI (2000) The yeast proteome database (YPD) and *Caenorhabditis elegans* proteome database (WormPD): comprehensive resources for the organization and comparison of model organism protein information. Nucleic Acids Res 28: 73–76

Craig RJ, Gallaher SD, Shu S, Salome P, Jenkins JW, Blaby-Haas CE, Purvine SO, O’Donnell S, Barry K, Grimwood J, Strenkert D, Kropat J, Daum C, Yoshinaga Y, Goodstein DM, Vallon O, Schmutz J, Merchant SS (2022) The Chlamydomonas Genome Project, version 6: reference assemblies for mating type *plus* and *minus* strains reveal extensive structural mutation in the laboratory. Plant Cell

Cross FR (2015) Tying Down Loose Ends in the *Chlamydomonas* Genome: Functional Significance of Abundant Upstream Open Reading Frames. G3 (Bethesda) 6: 435–446

Dall’Osto L, Bressan M, Bassi R (2015) Biogenesis of light harvesting proteins. Biochim Biophys Acta 1847: 861–871

Dobin A, Davis CA, Schlesinger F, Drenkow J, Zaleski C, Jha S, Batut P, Chaisson M, Gingeras TR (2013) STAR: ultrafast universal RNA-seq aligner. Bioinformatics 29: 15–21

Döring K, Ahmed N, Riemer T, Suresh HG, Vainshtein Y, Habich M, Riemer J, Mayer MP, O’Brien EP, Kramer G, Bukau B (2017) Profiling Ssb-Nascent Chain Interactions Reveals Principles of Hsp70-Assisted Folding. Cell 170: 298–311 e220

Eberhard S, Drapier D, Wollman FA (2002) Searching limiting steps in the expression of chloroplast-encoded proteins: relations between gene copy number, transcription, transcript abundance and translation rate in the chloroplast of *Chlamydomonas reinhardtii*. The Plant journal: for cell and molecular biology 31: 149–160

Floris M, Bassi R, Robaglia C, Alboresi A, Lanet E (2013) Post-transcriptional control of light-harvesting genes expression under light stress. Plant Mol Biol 82: 147–154

Forsythe ES, Grover CE, Miller ER, Conover JL, Arick MA, 2nd, Chavarro MCF, Leal-Bertioli SCM, Peterson DG, Sharbrough J, Wendel JF, Sloan DB (2022) Organellar transcripts dominate the cellular mRNA pool across plants of varying ploidy levels. Proc Natl Acad Sci U S A 119: e2204187119

Frank J, Agrawal RK (2000) A ratchet-like inter-subunit reorganization of the ribosome during translocation. Nature 406: 318–322

Fu Y, Wu PH, Beane T, Zamore PD, Weng Z (2018) Elimination of PCR duplicates in RNA-seq and small RNA-seq using unique molecular identifiers. BMC Genomics 19: 531

Fujita T, Kurihara Y, Iwasaki S (2019) The Plant Translatome Surveyed by Ribosome Profiling. Plant Cell Physiol 60: 1917–1926

Gallaher SD, Fitz-Gibbon ST, Strenkert D, Purvine SO, Pellegrini M, Merchant SS (2018) High-throughput sequencing of the chloroplast and mitochondrion of *Chlamydomonas reinhardtii* to generate improved de novo assemblies, analyze expression patterns and transcript speciation, and evaluate diversity among laboratory strains and wild isolates. Plant J 93: 545–565

Gao Y, Thiele W, Saleh O, Scossa F, Arabi F, Zhang H, Sampathkumar A, Kuhn K, Fernie A, Bock R, Schottler MA, Zoschke R (2022) Chloroplast translational regulation uncovers nonessential photosynthesis genes as key players in plant cold acclimation. Plant Cell 34: 2056–2079

Gawronski P, Jensen PE, Karpinski S, Leister D, Scharff LB (2018) Plastid ribosome pausing is induced by multiple features and is linked to protein complex assembly. Plant Physiol

Gerashchenko MV, Gladyshev VN (2014) Translation inhibitors cause abnormalities in ribosome profiling experiments. Nucleic Acids Res 42: e134

Gerashchenko MV, Gladyshev VN (2017) Ribonuclease selection for ribosome profiling. Nucleic Acids Res 45: e6

Gloge F, Becker AH, Kramer G, Bukau B (2014) Co-translational mechanisms of protein maturation. Curr Opin Struct Biol 24: 24–33

Harris CR, Millman KJ, van der Walt SJ, Gommers R, Virtanen P, Cournapeau D, Wieser E, Taylor J, Berg S, Smith NJ, Kern R, Picus M, Hoyer S, van Kerkwijk MH, Brett M, Haldane A, Del Rio JF, Wiebe M, Peterson P, Gerard-Marchant P, Sheppard K, Reddy T, Weckesser W, Abbasi H, Gohlke C, Oliphant TE (2020) Array programming with NumPy. Nature 585: 357–362

Hinnebusch AG (2011) Molecular mechanism of scanning and start codon selection in eukaryotes. Microbiol Mol Biol Rev 75: 434–467, first page of table of contents

Hsu PY, Calviello L, Wu HL, Li FW, Rothfels CJ, Ohler U, Benfey PN (2016) Super-resolution ribosome profiling reveals unannotated translation events in Arabidopsis. Proc Natl Acad Sci U S A

Hunter J (2007) Matplotlib: A 2D Graphics Environment. Computing in Science & Engineering 9: 90–95

Ingolia NT (2014) Ribosome profiling: new views of translation, from single codons to genome scale. Nat Rev Genet 15: 205–213

Ingolia NT (2016) Ribosome Footprint Profiling of Translation throughout the Genome. Cell 165: 22–33

Ingolia NT, Brar GA, Rouskin S, McGeachy AM, Weissman JS (2012) The ribosome profiling strategy for monitoring translation in vivo by deep sequencing of ribosome-protected mRNA fragments. Nat Protoc 7: 1534–1550

Ingolia NT, Ghaemmaghami S, Newman JR, Weissman JS (2009) Genome-wide analysis in vivo of translation with nucleotide resolution using ribosome profiling. Science 324: 218–223

Ingolia NT, Lareau LF, Weissman JS (2011) Ribosome profiling of mouse embryonic stem cells reveals the complexity and dynamics of mammalian proteomes. Cell 147: 789–802

Juntawong P, Girke T, Bazin J, Bailey-Serres J (2014) Translational dynamics revealed by genome-wide profiling of ribosome footprints in *Arabidopsis*. Proc Natl Acad Sci U S A 111: E203–212

Klimmek F, Sjodin A, Noutsos C, Leister D, Jansson S (2006) Abundantly and rarely expressed *Lhc* protein genes exhibit distinct regulation patterns in plants. Plant Physiol 140: 793–804

Komine Y, Kwong L, Anguera MC, Schuster G, Stern DB (2000) Polyadenylation of three classes of chloroplast RNA in *Chlamydomonas reinhadtii*. RNA 6: 598–607

Kramer G, Shiber A, Bukau B (2019) Mechanisms of Cotranslational Maturation of Newly Synthesized Proteins. Annu Rev Biochem 88: 337–364

Kropat J, Hong-Hermesdorf A, Casero D, Ent P, Castruita M, Pellegrini M, Merchant SS, Malasarn D (2011) A revised mineral nutrient supplement increases biomass and growth rate in *Chlamydomonas reinhardtii*. Plant J 66: 770–780

Lareau LF, Hite DH, Hogan GJ, Brown PO (2014) Distinct stages of the translation elongation cycle revealed by sequencing ribosome-protected mRNA fragments. Elife 3: e01257

Lauria F, Tebaldi T, Bernabo P, Groen EJN, Gillingwater TH, Viero G (2018) riboWaltz: Optimization of ribosome P-site positioning in ribosome profiling data. PLoS Comput Biol 14: e1006169

Lei L, Shi J, Chen J, Zhang M, Sun S, Xie S, Li X, Zeng B, Peng L, Hauck A, Zhao H, Song W, Fan Z, Lai J (2015) Ribosome profiling reveals dynamic translational landscape in maize seedlings under drought stress. Plant J 84: 1206–1218

Li H, Handsaker B, Wysoker A, Fennell T, Ruan J, Homer N, Marth G, Abecasis G, Durbin R, Genome Project Data Processing S (2009) The Sequence Alignment/Map format and SAMtools. Bioinformatics 25: 2078–2079

Liu MJ, Wu SH, Wu JF, Lin WD, Wu YC, Tsai TY, Tsai HL, Wu SH (2013) Translational landscape of photomorphogenic *Arabidopsis*. Plant Cell 25: 3699–3710

Lukoszek R, Feist P, Ignatova Z (2016) Insights into the adaptive response of *Arabidopsis thaliana* to prolonged thermal stress by ribosomal profiling and RNA-Seq. BMC Plant Biol 16: 221

Martin M (2011) Cutadapt removes adapter sequences from high-throughput sequencing reads. 2011 17: 3

McKinney W (2010) Data Structures for Statistical Computing in Python. In Proceedings of the 9th Python in Science Conference, pp 56–61

Merchante C, Brumos J, Yun J, Hu Q, Spencer KR, Enriquez P, Binder BM, Heber S, Stepanova AN, Alonso JM (2015) Gene-specific translation regulation mediated by the hormone-signaling molecule EIN2. Cell 163: 684–697

Moulin M, Nguyen GT, Scaife MA, Smith AG, Fitzpatrick TB (2013) Analysis of *Chlamydomonas* thiamin metabolism in vivo reveals riboswitch plasticity. Proc Natl Acad Sci U S A 110: 14622–14627

Mussgnug JH, Wobbe L, Elles I, Claus C, Hamilton M, Fink A, Kahmann U, Kapazoglou A, Mullineaux CW, Hippler M, Nickelsen J, Nixon PJ, Kruse O (2005) NAB1 is an RNA binding protein involved in the light-regulated differential expression of the light-harvesting antenna of *Chlamydomonas reinhardtii*. Plant Cell 17: 3409–3421

Nickelsen J, Bohne A-V, Westhoff P (2014) Chloroplast gene expression - translation. 49–78

Payne SH (2015) The utility of protein and mRNA correlation. Trends Biochem Sci 40: 1–3

Pechmann S, Willmund F, Frydman J (2013) The ribosome as a hub for protein quality control. Mol Cell 49: 411–421

Plancke C, Vigeolas H, Hohner R, Roberty S, Emonds-Alt B, Larosa V, Willamme R, Duby F, Onga Dhali D, Thonart P, Hiligsmann S, Franck F, Eppe G, Cardol P, Hippler M, Remacle C (2014) Lack of isocitrate lyase in *Chlamydomonas* leads to changes in carbon metabolism and in the response to oxidative stress under mixotrophic growth. Plant J 77: 404–417

Rooijers K, Loayza-Puch F, Nijtmans LG, Agami R (2013) Ribosome profiling reveals features of normal and disease-associated mitochondrial translation. Nat Commun 4: 2886

Russell JB, Cook GM (1995) Energetics of bacterial growth: balance of anabolic and catabolic reactions. Microbiol Rev 59: 48–62

Schneider-Poetsch T, Ju J, Eyler DE, Dang Y, Bhat S, Merrick WC, Green R, Shen B, Liu JO (2010) Inhibition of eukaryotic translation elongation by cycloheximide and lactimidomycin. Nat Chem Biol 6: 209–217

Schroda M, Hemme D, Mühlhaus T (2015) The *Chlamydomonas* heat stress response. Plant J 82: 466–480

Schuster M, Gao Y, Schöttler MA, Bock R, Zoschke R (2019) Limited Responsiveness of Chloroplast Gene Expression during Acclimation to High Light in Tobacco. Plant Physiol

Sharma P, Wu J, Nilges BS, Leidel SA (2021) Humans and other commonly used model organisms are resistant to cycloheximide-mediated biases in ribosome profiling experiments. Nat Commun 12: 5094

Slobodin B, Dikstein R (2020) So close, no matter how far: multiple paths connecting transcription to mRNA translation in eukaryotes. EMBO Rep 21: e50799

Smith T, Heger A, Sudbery I (2017) UMI-tools: modeling sequencing errors in Unique Molecular Identifiers to improve quantification accuracy. Genome Res 27: 491–499

Sun Y, Zerges W (2015) Translational regulation in chloroplasts for development and homeostasis. Biochim Biophys Acta 1847: 809–820

Teixeira FK, Lehmann R (2019) Translational Control during Developmental Transitions. Cold Spring Harb Perspect Biol 11

Ting MKY, Gao Y, Barahimipour R, Ghandour R, Liu J, Martinez-Seidel F, Smirnova J, Gotsmann VL, Fischer A, Haydon MJ, Willmund F, Zoschke R (2023) Improved Plant Ribosome Profiling with Structural Assessment of rRNA Contaminants. manuscript submitted

Trösch R, Barahimipour R, Gao Y, Badillo-Corona JA, Gotsmann VL, Zimmer D, Mühlhaus T, Zoschke R, Willmund F (2018) Commonalities and differences of chloroplast translation in a green alga and land plants. Nat Plants 4: 564–575

Trösch R, Ries F, Westrich LD, Gao Y, Herkt C, Hoppstädter J, Heck-Roth J, Mustas M, Scheuring D, Choquet Y, Räschle M, Zoschke R, Willmund F (2022) Fast and global reorganization of the chloroplast protein biogenesis network during heat acclimation. Plant Cell 34: 1075–1099

Virtanen P, Gommers R, Oliphant TE, Haberland M, Reddy T, Cournapeau D, Burovski E, Peterson P, Weckesser W, Bright J, van der Walt SJ, Brett M, Wilson J, Millman KJ, Mayorov N, Nelson ARJ, Jones E, Kern R, Larson E, Carey CJ, Polat I, Feng Y, Moore EW, VanderPlas J, Laxalde D, Perktold J, Cimrman R, Henriksen I, Quintero EA, Harris CR, Archibald AM, Ribeiro AH, Pedregosa F, van Mulbregt P, SciPy C (2020) SciPy 1.0: fundamental algorithms for scientific computing in Python. Nat Methods 17: 261–272

Wang F, Dischinger K, Westrich LD, Meindl I, Egidi F, Trosch R, Sommer F, Johnson X, Schroda M, Nickelsen J, Willmund F, Vallon O, Bohne AV (2023) ONE-HELIX PROTEIN 2 is not required for the synthesis of photosystem II subunit D1 in Chlamydomonas. Plant Physiol

Wang R, Amoyel M (2022) mRNA Translation Is Dynamically Regulated to Instruct Stem Cell Fate. Front Mol Biosci 9: 863885

Waskom M (2021) Seaborn: statistical data visualization. Journal of Open Source Software 6: 3021

Westrich LD, Gotsmann VL, Herkt C, Ries F, Kazek T, Trosch R, Armbruster L, Mühlenbeck JS, Ramundo S, Nickelsen J, Finkemeier I, Wirtz M, Storchova Z, Räschle M, Willmund F (2021) The versatile interactome of chloroplast ribosomes revealed by affinity purification mass spectrometry. Nucleic Acids Res 49: 400–415

Wolin SL, Walter P (1988) Ribosome pausing and stacking during translation of a eukaryotic mRNA. EMBO J 7: 3559–3569

Wu CC, Zinshteyn B, Wehner KA, Green R (2019) High-Resolution Ribosome Profiling Defines Discrete Ribosome Elongation States and Translational Regulation during Cellular Stress. Mol Cell 73: 959–970 e955

Wu HL, Song G, Walley JW, Hsu PY (2019) The Tomato Translational Landscape Revealed by Transcriptome Assembly and Ribosome Profiling. Plant Physiol 181: 367–380

Yang X, Cui J, Song B, Yu Y, Mo B, Liu L (2020) Construction of High-Quality Rice Ribosome Footprint Library. Front Plant Sci 11: 572237

Yang X, Song B, Cui J, Wang L, Wang S, Luo L, Gao L, Mo B, Yu Y, Liu L (2021) Comparative ribosome profiling reveals distinct translational landscapes of salt-sensitive and -tolerant rice. BMC Genomics 22: 612

Zhang W, Dunkle JA, Cate JH (2009) Structures of the ribosome in intermediate states of ratcheting. Science 325: 1014–1017

Zoschke R, Bock R (2018) Chloroplast Translation: Structural and Functional Organization, Operational Control and Regulation. Plant Cell 30: 745–770

